# A syntenin-deficient microenvironment educates AML for aggressiveness

**DOI:** 10.1101/2021.01.06.425538

**Authors:** R Leblanc, J Fares, A Goubard, R Castellano, L Camoin, M Balzano, R Ghossoub, B Bou-Tayeh, C Fauriat, N Vey, JP Borg, Y Collette, M Aurrand-Lions, G David, P Zimmermann

## Abstract

In acute myeloid leukemia (AML), the stromal microenvironment plays a prominent role in promoting tumor cell survival and progression. Although widely explored, the crosstalk between leukemic and stromal cells remains poorly understood. Syntenin, a multi-domain PDZ protein, controls both the trafficking and signaling of key molecules involved in intercellular communication. Therefore, we aimed to clarify the role of environmental syntenin in the progression of AML. By *in vivo* approaches in syngeneic mice, we demonstrate that a syntenin-deficient environment reprograms AML blasts to survive independently of the stroma. Up-regulation of EEF1A2 in the blasts controls this gain of cell survival. Furthermore, using *ex vivo* co-culture systems, we show that syntenin-deficient bone marrow stromal cells (BMSC) enhance the survival of different types of AML cells, including patient samples, and suffice to educate syngeneic AML, recapitulating micro-environmental effects observed *in vivo.* We establish that syntenin-deficiency causes an increase of eIF5A and autophagy-related factors in BMSC, and provide evidence that the inhibition of autophagy prevents syntenin-deficient BMSC to stimulate AML survival. Altogether, these findings indicate that host-syntenin in the BM microenvironment acts as a repressor of AML aggressiveness.

**Key points:**

- A syntenin-deficient host reprograms AML blasts, enhancing total protein synthesis and cell survival pathways
- Autophagy in the syntenin-deficient microenvironment is responsible for the gain of AML cell survival

## Introduction

Acute myeloid leukemia (AML) accounts for over 80% of all acute leukemias in adults with a five-year overall survival rate below 30%^1^. AML is characterized by clonal expansion of hematopoietic cells at distinctive stages of differentiation and a block in hematopoietic differentiation. At late stage of the disease, the bone marrow (BM) can no longer ensure the production of normal blood cells.

Hematopoietic development is regulated by bone marrow stromal cells (BMSC), which establish and sustain communication pathways needed to maintain hematopoietic homeostasis. Yet, within a leukemic niche, the BM microenvironment that hosts the leukemia appears corrupted, promoting the growth and survival of leukemic cells^2–4^. BMSC-mediated protection of leukemic cells relies on leuko-stromal interactions mediated by adhesion molecules, cytokines, chemokines, growth factors and cognate receptors. In addition, the release of extracellular vesicles produced by non-haemopoietic BM cells and mitochondrial transfers have been recognized as important mechanisms of cellular crosstalk between leukemic cells and surrounding BM microenvironment^5^. Recent studies also implicate autophagy in the tumor stroma, supplying metabolites to tumor cells, in leukemia progression and survival^6^. Thus, better understanding of molecular mechanisms supporting non-haemopoietic BM signaling to leukemic cells can potentially help to refine anti-cancer therapies.

Syntenin is an evolutionary conserved and ubiquitously expressed PDZ scaffold protein interacting with various transmembrane receptors directly and/or with the help of syndecan heparan sulfate proteoglycans^7–9^. It impacts on their vesicular trafficking^10^, affecting multiple processes, including immune cell regulation, neuronal differentiation and anti-viral activity^11^. Noteworthy, syntenin controls the formation of endosomal intraluminal vesicles and the secretion of exosomes^12,13^, as well as regulates the uptake of exosomes^14^. Regulating exosome formation and release, syntenin potentially also provides pathways for unconventional protein secretion, including that of several cytokines and co-stimulatory molecules^15^. Syntenin might thus have an important role in leukemia progression by regulating intercellular communications within the leukemic niche.

Gain of syntenin expression in tumor cells has been unambiguously associated with the invasion and the metastatic potential of various solid cancers including melanoma, glioblastoma, prostate and head/neck squamous cancers^16, 17^,^18^,^19^ and identified as regulator of protective autophagy^20^. Syntenin has therefore been proposed as a potentially valuable target for cancer therapy^21^. In comparison, the impact on tumor development resulting from syntenin suppression in the tumor microenvironment remains under-explored. Syntenin-deficiency of the host was shown to reduce melanoma metastasis, by dampening tumor-supporting inflammation^22^.

Considering the pathophysiological importance of the microenvironment in AML, we investigated the impact of loss of syntenin expression *in vivo* and *ex vivo,* in various experimental models of AML. Our observations contrast with the data reported in melanoma, showing that bone marrow microenvironmental syntenin acts as negative regulator of AML progression, raising concerns and potentially cautions regarding systemic syntenin-targeting in AML therapy.

## Methods

### **Antibodies and reagents** are listed in **Table S1**

#### Cell lines and patient samples

Human AML cell lines U937 and HL60 were purchased from the American Type culture collection (ATCC; Manassas, USA) and maintained in minimum essential medium-α medium (Gibco, Carlsbad, USA). Murine AML cell line C1498 was obtained from ATCC and cultured in Dulbecco’s modified eagle’s medium (Gibco, Carlsbad, USA). All cell lines were grown in media supplemented with 10% FBS (Eurobio, Les Ulis, France) and incubated at 37 °C, with 5% CO2. The cultured cells were split every 2-3 days, maintaining an exponential growth. Syngeneic FLB1 model is described elsewhere^23^. Patient samples were obtained from the IPC/CRCM Tumor Bank / Biological Resource Center for Oncology that operates under authorization # AC-2018-1905 granted by the French Ministry of Research. Prior to scientific use of samples and data, patients were appropriately informed and asked for consent in writing, in compliance with French and European regulations. This study was performed after approval by our institutional review board.

#### Mice

C57BL/6J mice were purchased from Janvier Laboratories, France. Both male and female mice were used between 6 and 11 weeks of age and were housed under specific pathogen-free conditions. Syntenin-knockout animals were generated as previously described^14^ and the hygromycin-resistance gene was removed using Flipper mice^24^.

#### Animals studies

All experiments were performed in compliance with the laws and protocols approved by animal ethics committees (Agreement No. APAFIS#5123-2016041809182209v2). For *in vivo* expansion, FLB1 cells were injected into the retro-orbital vein. Leukemia progression was monitored weekly by FACS analysis using CD45.1-FITC and CD45.2-V450 antibodies. Recipient mice were sacrificed when blast levels reached >10% in the PB (survival threshold) and leukemic cells were collected from femurs and tibias. For serial transplantation assays, blasts collected from the BM (>90% blast invasion) of 3 different animals were pooled and used (5×10^4^ cells/mouse) for re-injection in the following transplantation round.

#### Co-culture experiments

Passage 4 to 10 BMSC WT/KO were detached, counted and irradiated at 30Gy. A total of 10,000 irradiated BMSC were seeded in 12-well plates pre-coated with type-I collagen (5μg/cm^2^; Gibco) and were cultured in complete medium for 24h. Then, 50,000 AML cells were added to each well and maintained in short-term co-culture. HL60 and FLB1 cell lines were also maintained in long-term co-culture for 1 month, with a media change every 3-4 days.

#### AML cell viability

The possible role of exosomes was investigated by using conditioned media, isolated after 24h from both WT and KO irradiated BMSC (30Gy) and depleted or not depleted of exosomes (by ultracentrifugation; 100,00g at 4°C for 1h). After 48 to 96h of co-culture/treatment, AML blasts were collected, and early/late apoptosis was measured by using the PE Annexin V apoptosis detection kit with 7-AAD (Biolegend, San Diego, USA) according to the manufacturer. Annexin V positive but 7-AAD negative (early apoptotic cells) and both Annexin V and 7-AAD positive cells (late-stage apoptosis) were determined by using FACS LSRII flow cytometer and data were analyzed with Flowjo software (BD Biosciences, Franklin Lakes, USA).

#### Mass Spectrometry Analysis

FLB1 cells, isolated after 4 rounds of serial transplantation from five WT and KO mice for each condition, were sorted by flow cytometry (BDFACs ARIA III) using CD45.1 and CD45.2 antibodies. Two million of CD45.1+ cells were lysed in RIPA buffer and analyzed by mass spectrometry as described in supplementary methods.

#### Statistical analysis

Differences between groups were determined by 1-way or 2-way analysis of variance (ANOVA), followed by a Bonferroni posttest using GraphPad Prism v8.0c software. Single comparisons were carried out using the parametric Student t-test or the nonparametric Mann-Whitney U test. P < .05 was considered statistically significant.

## Results

### Host-syntenin deficiency supports AML aggressiveness

To address the role of host-syntenin in AML development, we used the FBL1 murine AML model, created through retroviral overexpression of AML oncogenes (i.e. Meis1 and Hoxa9) in C57BL/6 fetal liver cells. This model has a frequency of leukemic stem cells that is comparable to that of the human disease and remains stable over successive transplantations^23^ (**Figure 1A & 1B**). In non-irradiated WT recipient mice, FLB1 cells engraft successfully and produce widespread AML within three weeks. At graft 1, loss of host-syntenin had no noticeable effect on FLB1 progression (**Figure S1A**). In parallel short-term homing assays, lack of host-syntenin had also no significant effect on the recruitment of the leukemic cells to the primary and secondary hematopoietic organs (**Figure S1B**). Yet in serial transplantation experiments, from graft 3 on, the AML becomes more aggressive in a syntenin-negative environment (**Figure 1B**). Serial back-transplantation experiments (**Figure 1A**) did not revert leukemic progression, supporting the notion that the blasts acquired a cell-autonomous aggressiveness (**Figure 1C**).

**Figure 1.**
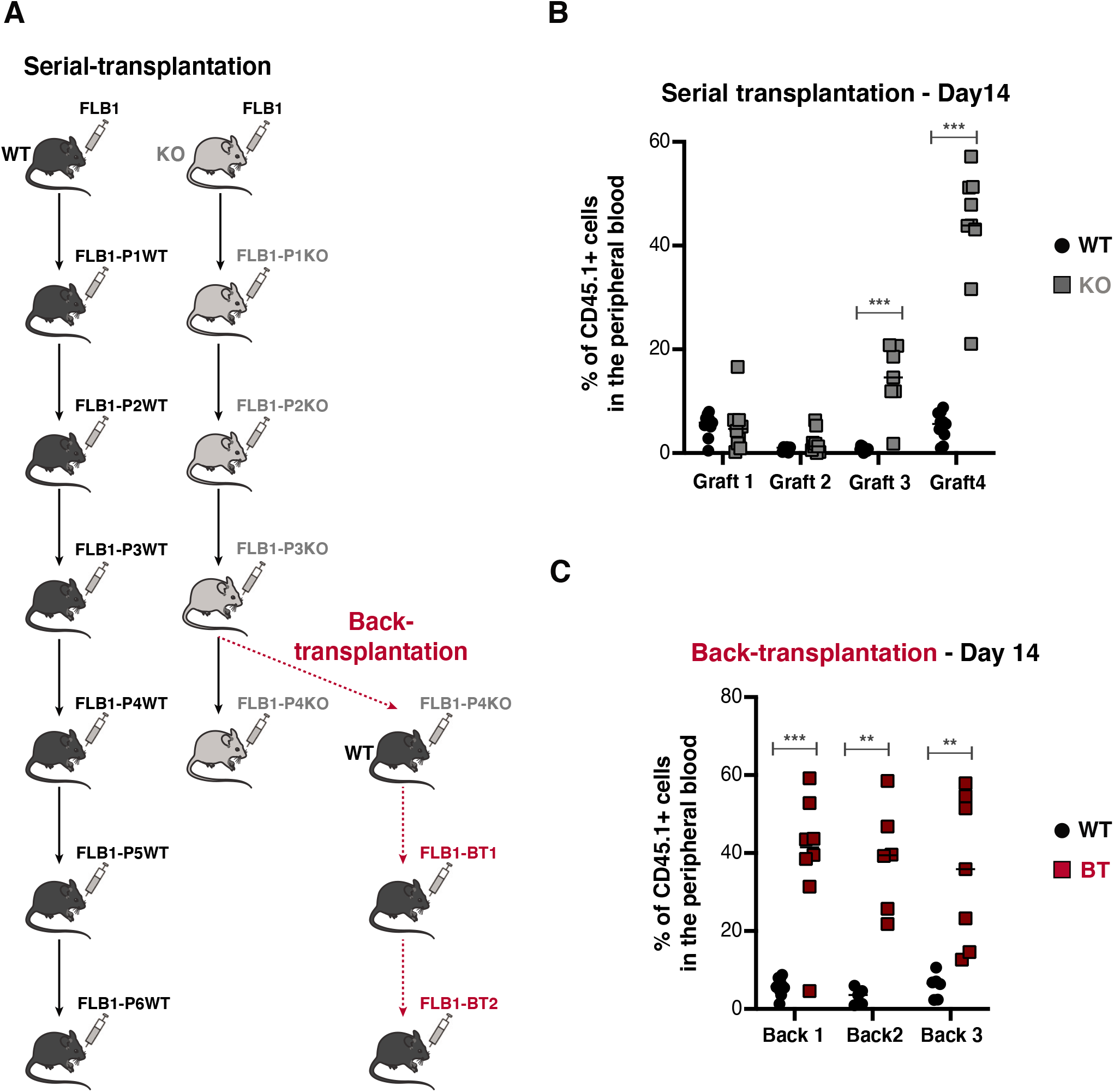
FLB1 blasts acquire cell-autonomous aggressiveness upon serial transplantation in a syntenin-deficient host. (A) WT and KO mice were injected with FLB1 and sacrificed when blast levels reached >10% in the PB (survival threshold). Blasts collected from the BM (>90% blast invasion, see **Figure S1C & 1D**) of 3 different animals were pooled and used for re-injection in the following transplantation or back-transplantation round. (B) Blood samples were collected from mice on day 14 post injection for FACS analysis. Data from 4 sequential grafts/passages (P1-P4) are shown here. Results are expressed as the percentage of CD45.1^+^ cells in the PB, ± SEM. Statistical analysis was performed using the nonparametric Mann-Whitney U test (***, P < 0.0001). (C) FLB1 having undergone 4 serial transplantations in KO animals were then re-transplanted in WT animals. Blood samples were collected from mice on day 14 post injection for FACS analysis. Data from 3 sequential back-transplantation are shown here (Back 1 to Back 3). Results are expressed as percentage of CD45.1^+^ cells in the PB ± SEM. Statistical analysis was performed using the nonparametric Mann-Whitney U test (**, P<0.001***, P < 0.0001).

### The syntenin-deficient host reprograms AML blasts, enhancing total protein synthesis and cell survival pathways

To elucidate the mechanisms underlying the gain of aggressiveness, we first assessed “basic” pathways involved in leukemia progression. As shown in **Figure S2**, we excluded altered homing, cell proliferation or change in leukemic initiating cell frequencies. We then compared the proteomes of sorted FLB1 cells, isolated from WT and KO mice after 4 rounds of serial transplantation (designated as FLB1-P4WT and FLB1-P4KO). Quantitative protein expression analysis by mass spectrometry allowed the reproducible identification of 1,961 proteins differentially expressed between recovered FLB1 cells (**Table S2**). **Figure 2A** shows a volcano plot of the entire data set, highlighting proteins whose expressions were significantly different within FLB1-P4WT and -P4KO cells. The selected thresholds revealed 62 up-regulated (blue dots) and 69 down-regulated (orange dots) proteins in FLB1 isolated from KO animals. Ingenuity pathway analysis of these protein changes (**Table S3**) predicted the activation of functions implicated in “protein synthesis” and the inhibition of functions related to “cell death” in FLB1-P4KO cells (**Figure 2B**).

**Figure 2.**
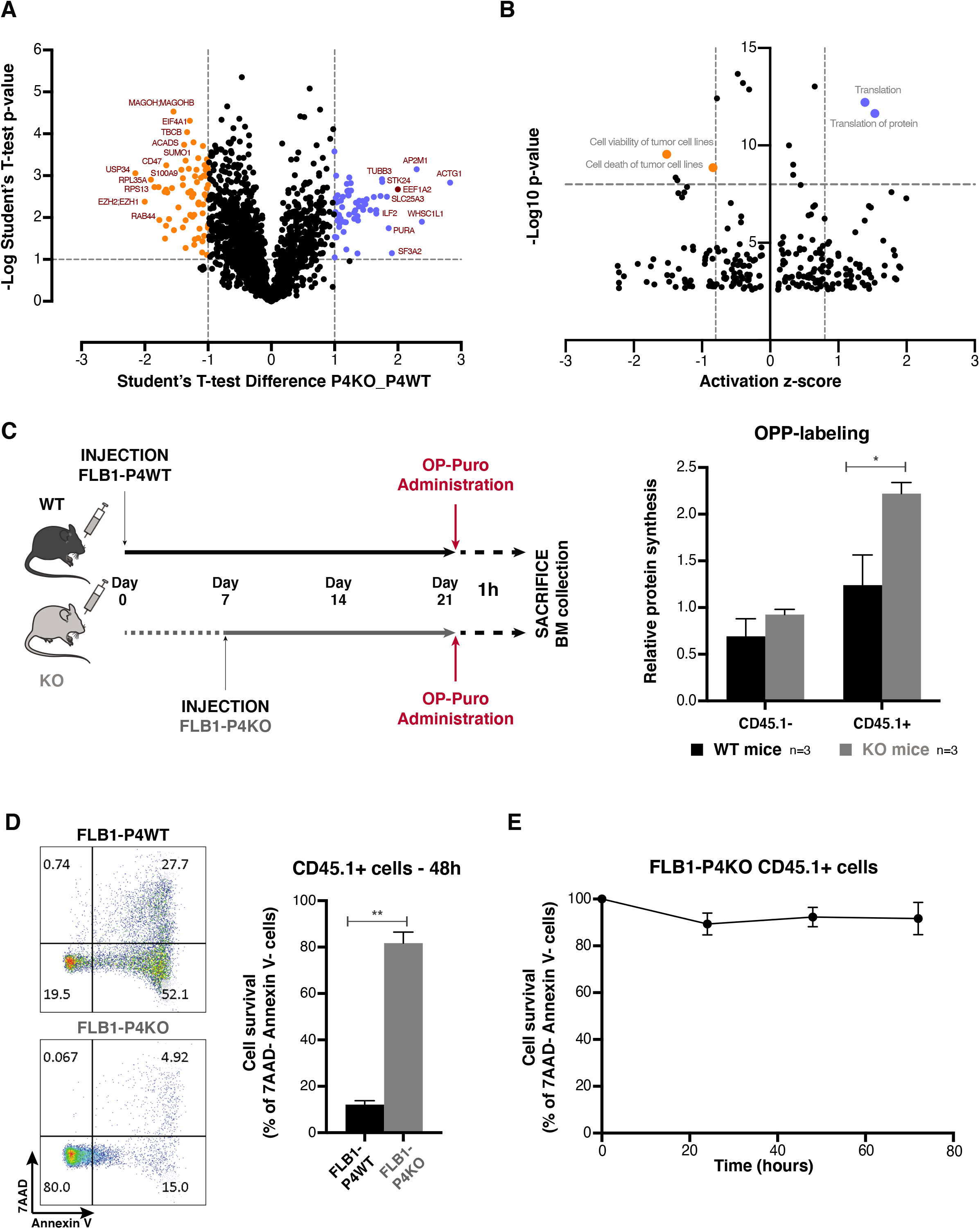
Education by a syntenin-deficient host sustains total protein synthesis and cell survival in blast cells. (A) Volcano plots showing differential protein expressions, with log10 levels (x-axis) and -log(p-value) (y-axis), in sorted FLB1-P4WT and FLB1-P4KO blasts (see **Table S2**). (B) Dot plot representing the corresponding activation of different categories of biological function, based on ‘Diseases & Functions’ annotation, with the activation z-score (x- axis) and the -log10(p-value) (y-axis). (C) FLB1-P4WT cells were injected at day 0 into WT mice, while the inoculation of FLB1-P4KO cells into KO mice was delayed for one week. At similar stages of disease progression in the different hosts, when reaching similar tumor loads in the different animals, O-propargyl-puromycin (OP-Puro; 50mg/kg body mass) was injected intraperitoneally. One hour later BM was collected for total protein synthesis analysis. OPP incorporated into nascent polypeptide chains was then fluorescently labeled via “Click-it Chemistry” *(Left panel).* Graph representing the relative levels of protein synthesis in CD45.1^-^ and CD45.1^+^ cells, normalized to unfractionated BM cells ± SEM. Statistical analysis was performed using the nonparametric Mann-Whitney U test (*, P<0.05) *(Rightpanel).* (D) CD45.1^+^ blasts harvested from WT/KO animals at graft 4 were seeded in complete RPMI media for apoptosis assays. Results are represented as mean value of living (annexing, 7AAD^-^) CD45.1^+^ cells ± SEM performed in 3 independent experiments. Statistical analysis was performed using the nonparametric Mann-Whitney U test (**, P<0.001). (E) FLB1 blasts isolated from KO animals at graft 4 (FLB1-P4KO) were seeded in complete RPMI media and tested for apoptosis over the time. Results are represented as mean value of living (annexinV-, 7AAD^-^) CD45.1^+^ cells ± SEM, calculated from 3 independent experiments.

To validate these predictions, we first tested whether total protein synthesis was altered in FLB1 cells after 4 successive passages into syntenin-KO mice, using O-propargyl-puromycin labelling methods^25^(**Figure 2C**). Of note, total protein synthesis in the nonmalignant CD45.1^-^ host cells was similar in WT and KO mice (**Figure 2C & S3A**). In contrast, we noticed a significant increase in total protein synthesis in FLB1 maintained in syntenin-KO mice (**Figure 2C & S3A**). Secondly, to test for survival, CD45.1^+^ FLB1 cells were isolated from FLB1-P4WT and -P4KO BM aspirates by FACS and seeded in complete RPMI medium. While FLB1-P4WT show high apoptosis levels and fail to survive *ex vivo,* FLB1-P4KO cells display markedly reduced apoptosis following 48h *in vitro* culture (**Figure 2D**) with a survival rate that remained stable over time (**Figure 2E**), indicating the acquisition of cell autonomy and independence regarding the environment.

Altogether, our data reveal (i) a reprogramming of FLB1 cells upon serial transplantation into a syntenin-negative environment and (ii) a gain of aggressiveness seemingly related to enhanced protein synthesis associated with a cell survival advantage.

### EEF1A2 plays a pivotal role in the survival of AML cells educated in a syntenin deficient host

Because EEF1A2 was ranked as top upregulated factor in FLB1-P4KO cells (**Figure 2A**) and since it has anti-apoptotic functions through the activation of Akt/mTORC1/RPS6 signaling^26–28^, we examined the status of this pathway. Western blot analysis validated the mass spectrometry data, revealing a significant 6-fold increase of the EEF1A2 cellular levels (**Figure 3A**). FLB1-P4KO cells showed an increase of the panAKT and RPS6 cellular levels, respectively associated with an increased phosphorylation of pS473-AKT, pT450-AKT and pS235/236-RPS6 (**Figure 3B**). As AKT is a substrate of mTORC2 complex, we also investigated whether RICTOR contributes to AKT/RPS6 phosphorylation, but neither the cellular level nor pT1135 phosphorylation of RICTOR were affected (**Figure 3B**). We further investigated the relevance of EEF1A2 by treating FLB1-P4KO cells with increasing concentrations of metarrestin. Metarrestin is a perinucleolar compartment inhibitor, disrupting the nucleolar structure and inhibiting RNA polymerase I transcription and ribosome synthesis by interacting with the translation elongation factor EEF1A2^29^. A treatment for 12h, at concentrations increasing from 1.25 to 10μM, markedly reduced the cellular level of RPS6 in FLB1-P4KO, also decreasing pS473-AKT, pT450-AKT and pS235/236-RPS6 phosphorylation (**Figure 4A**). Metarrestin administration was also associated with a dose-dependent decrease of aggressive FLB1 survival *ex vivo*, at 24h and 72h of culture (**Figure 4B and 4C**). While admittedly already difficult to maintain *ex vivo,* FLB1-P4WT cells, in contrast, were not significantly affected by a 24h treatment with 10μM of metarrestin (**Figure 4C**).

**Figure 3.**
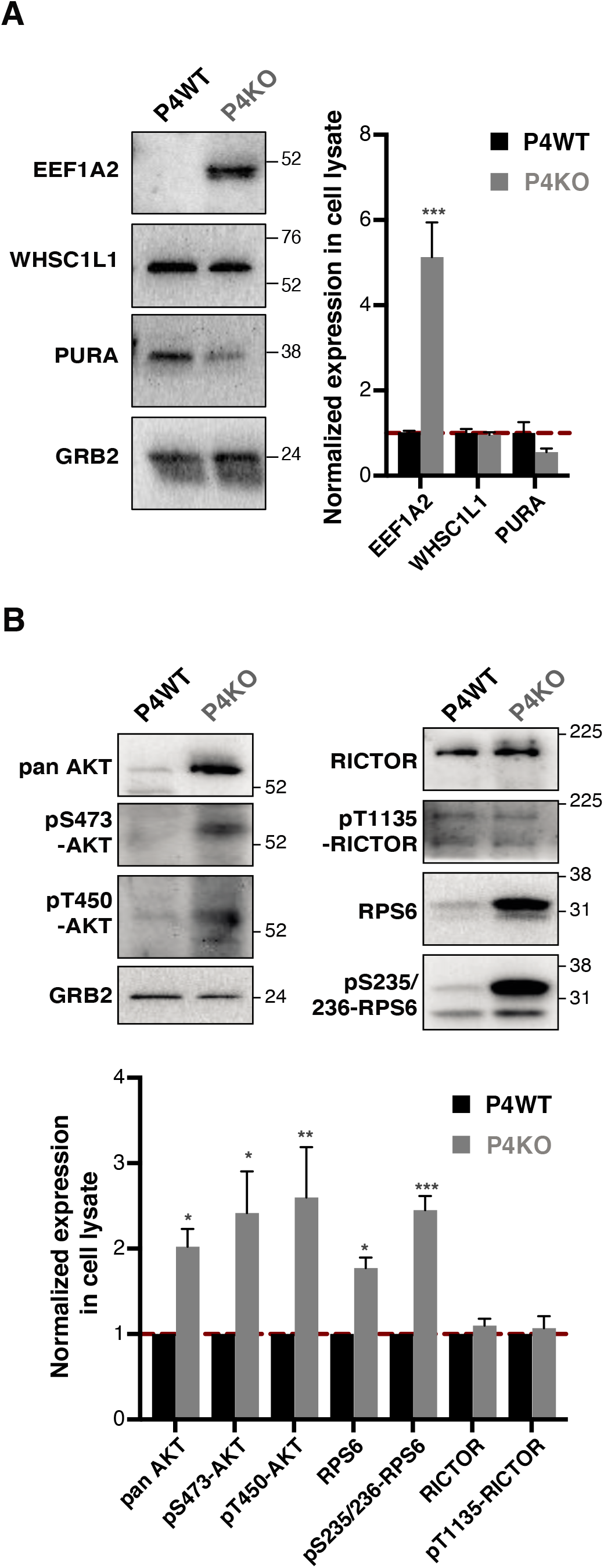
The EEF1A2 pathway is up-regulated in syntenin-KO-educated blasts. (A) & (B) Total cell lysates of P4WT and P4KO FLB1 were analyzed by western blot for different markers, as indicated. Histograms represent mean signal intensities ± SEM in cell lysates, relative to signals obtained from sorted FLB1-P4WT cells, calculated from the analysis of 5 independent mice of each category. Statistical analysis was performed using the one-way analysis of variance (ANOVA) (*P < 0.05; **P < 0.01; ***P < 0.001).

**Figure 4.**
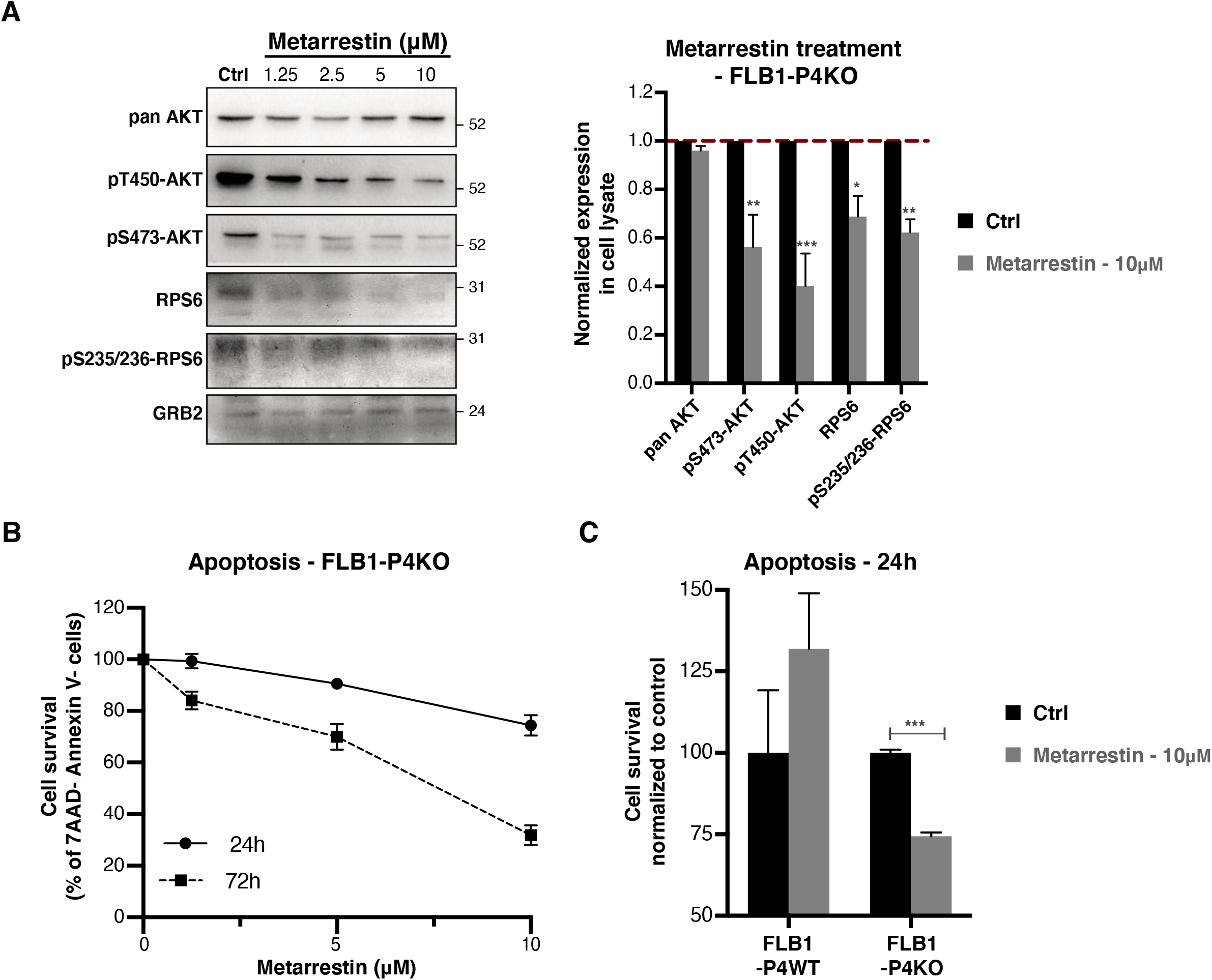
The EEF1A2 pathway supports the survival of syntenin-KO-educated blasts. (A) Aggressive FLB1-P4KO cells were treated with DMSO (control) or metarrestin (1.25 to 10μM) for 16h, in medium containing exosome-depleted FCS (10%). Total cell lysates were analyzed by western blot for different markers, as indicated. Histograms represent mean signal intensities ± SEM in cell lysates, relative to signals obtained from cells treated with DMSO (control), calculated from the analysis of 3 independent experiments. (B) Effect of metarrestin treatment on FLB1-P4KO cell apoptosis. Cell survival in the presence of increasing concentrations of metarrestin was evaluated at 24h and 72h. Data represent the mean percentage of living CD45.1^+^ cells ± SEM of 3 independent experiments performed in triplicate. (C) FLB1-P4WT and FLB1-P4KO were treated with DMSO (control) or 10μM of metarrestin for 24h. Data represent the percentage of viable (annexinV-, 7AAD^-^) cells normalized to control ± SEM, calculated from 3 independent experiments performed in triplicate. Statistical analysis was performed using the one-way analysis of variance (ANOVA) (***P < 0.001).

Altogether, our data suggest that gain of EEF1A2 plays a pivotal role in driving the gain of aggressiveness of FLB1 educated in syntenin-null mice.

### BMSC isolated from syntenin-KO mice convey AML gain of aggressiveness

Because BMSC were previously reported to support AML aggressiveness by diverse mechanisms^2,30,31^, we aimed to clarify whether syntenin-KO BMSC might suffice to support AML reprogramming. Validated primary BMSC isolated from both WT and KO animals (i.e. with mesenchymal pattern and differentiation ability; **Figure S4**), were co-injected with FLB1 cells in WT animals (**Figure 5A**). Co-injection of WT BMSC did not affect FLB1 progression, but peripheral blood levels and bone marrow invasion were significantly higher when FLB1 cells were injected together with KO BMSC (**Figure 5A**).

**Figure 5.**
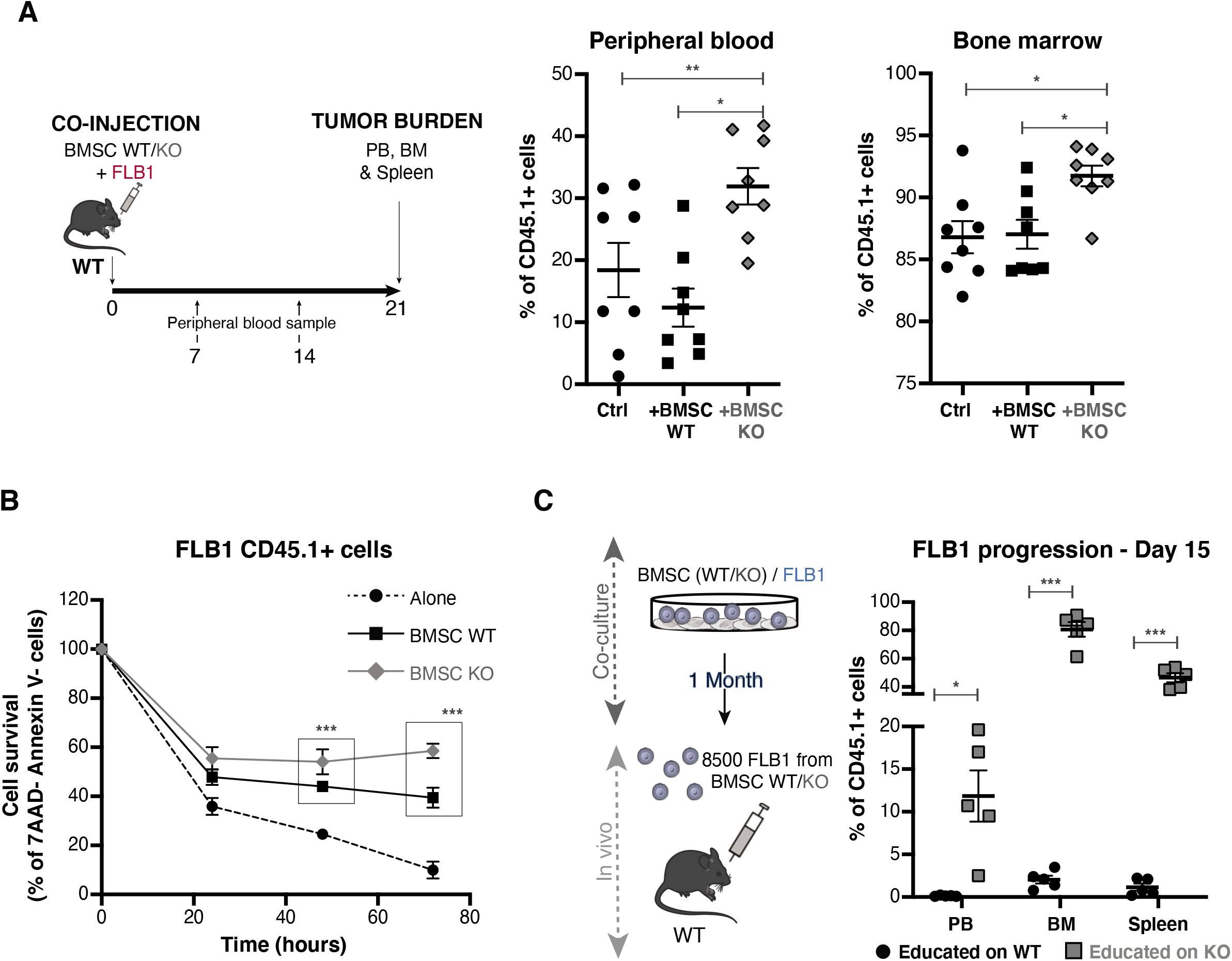
Syntenin-deficient BMSC enhance AML aggressiveness *in vivo.* (A) Non-irradiated 8-11w old WT mice were injected with 50,000 FLB1 cells ± 10,000 BMSC WT/KO in the retro-orbital vein. Leukemia progression was assessed weekly, by FACS analysis of a peripheral blood (PB) sample (Upper panel). Summary of the CD45.1^+^ cell frequencies measured in the PB, the BM and the spleen of the different mice in the different groups, 21 days after cell inoculation. Results are expressed as percentage of CD45.1^+^ cells, and calculated means ± SEM. Statistical analysis was performed using one-way analysis of variance (ANOVA) (*, P < 0.05; **, P < 0.01) (Lower panel). (B) FLB1 cells were cultured, alone or together with BMSC WT/KO, in medium containing exosome-depleted FCS (10%). Blasts were collected at the times indicated and stained for apoptosis markers. Results are expressed as mean percentages of living (AnnexinV^-^, 7AAD^-^) CD45.1^+^ cells, ± SEM, calculated from 3 independent experiments. Statistical analysis was performed using 1way ANOVA test (***, P < 0.001). (C) FLB1 cells were co-cultured with BMSC WT/KO in medium containing exosome-depleted FCS (10%). After one month, surviving FLB1 cells, ‘educated’ by BMSC WT/KO, were collected, counted and injected in the retro-orbital vein of WT mice (Left panel). Leukemia progression was assessed weekly, by FACS analysis. Summary of the CD45.1+ cell frequencies measured in PB, BM and the spleen of the different mice in the different groups, 14 days after cell inoculation. Results are expressed as percentage of CD45.1^+^ cells, and calculated means ± SEM. Statistical analysis was performed using 1way ANOVA test (*, P < 0.05; **, P < 0.005) (Right panel).

To further clarify whether BMSC might suffice to induce AML aggressiveness, we tested whether KO BMSCs would be competent to reprogram AML *in vitro.* In short-term co-cultures syntenin-deficient BMSC protected FLB1 cells from apoptosis over time (**Figure 5B**). In longterm co-culture experiments, i.e. after one-month, remaining CD45.1^+^ blasts were collected and injected (8,500 blasts per condition) in WT animals. Whereas flow cytometry analysis revealed an important invasion of the bone marrow and spleen by FLB1 co-cultured with KO BMSC, CD45.1+ cells were barely detectable in animals injected with FLB1 maintained on a wild-type stroma (**Figure 5C**).

Taken together, these results demonstrate that exposure to the stroma alone is able to recapitulate effects on AML cells observed *in vivo* in syntenin-null mice.

### BMSC with syntenin-KO increase the survival of various leukemia cell lines and AML blasts from patients

To evaluate whether the effects of syntenin-null BMSC are specific for the Hoxa9-Meis1 AML model or might be more generic, we also co-cultured established murine and human AML cell lines with WT or KO BMSC. Except for the C1498 murine cell line (with high spontaneous survival), loss of syntenin in BMSC significantly enhanced AML survival at 72h (**Figure 6A**). Similarly, primary human leukemic blasts, isolated from the peripheral blood or the bone marrow of 12 different AML patients and tested for apoptosis after 72h of co-culture with WT or KO BMSC, showed a significantly increased survival on a syntenin-deficient stroma (**Figure 6B & Table S4**). Gain of survival was primary sample-dependent, but subtype or mutationindependent (**Table S4**).

**Figure 6.**
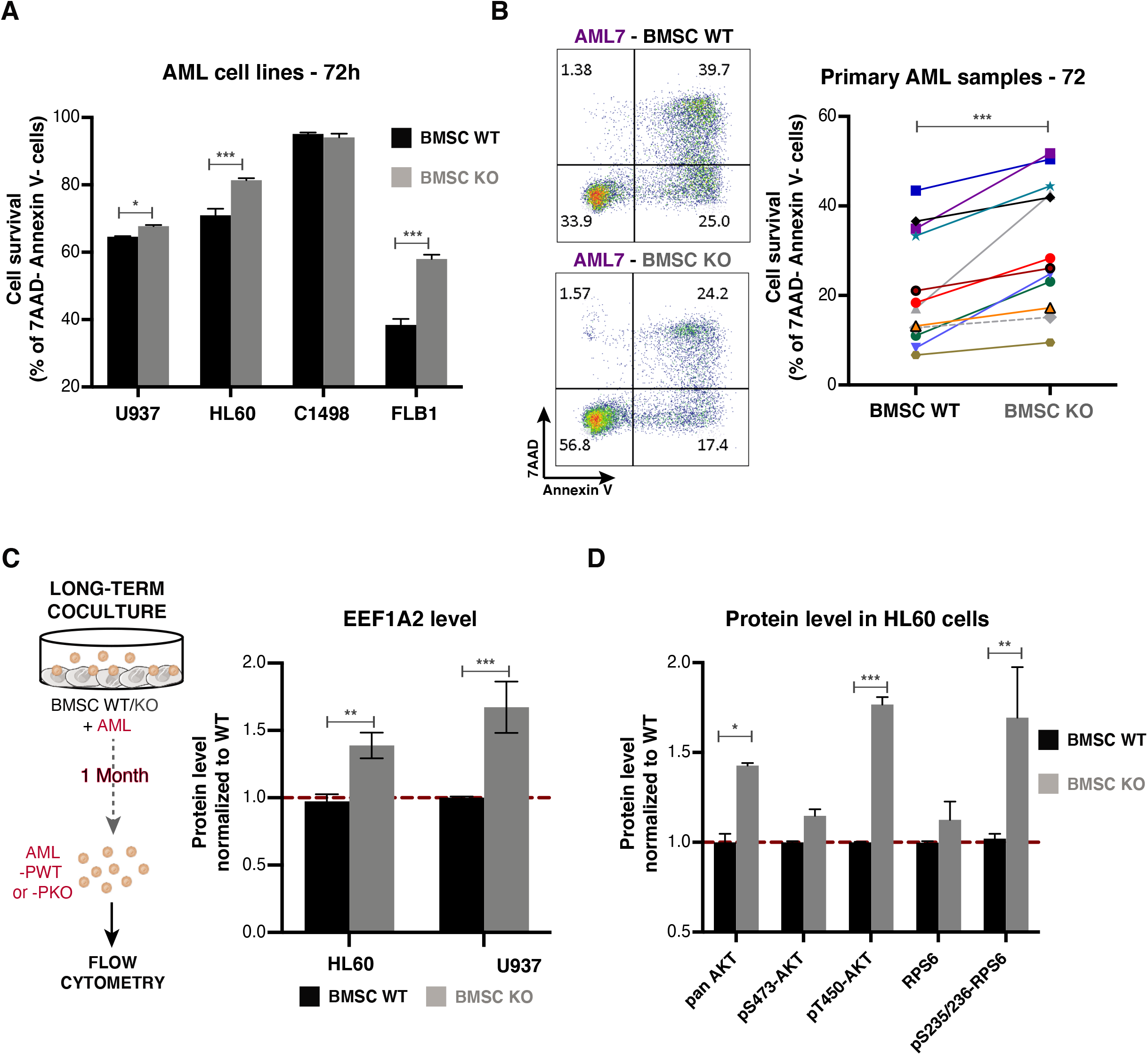
Syntenin loss in BMSC enhances the survival of patient samples and several cellular models of AML and ultimately reprograms AML blasts. (A) Human and murine AML cell lines were cultured in the presence of BMSC WT/KO in medium containing exosome- depleted FCS (10%). After 72h of co-culture, the AML cells were collected and stained for apoptosis markers. Results are expressed as mean percentages of living (AnnexinV^-^, 7AAD^-^) cells ± SEM, calculated from 3 independent experiments. Statistical analysis was performed using the two-way analysis of variance (ANOVA) (*, P < 0.05; ***, P < 0.01). (B) Twelve samples of different subtypes of primary AML were cultured in presence of BMSC WT/KO in medium containing exosome-depleted FCS (10%). After 72h of co-culture, the blasts were collected and stained for apoptosis markers. Results are expressed as mean percentages of living (AnnexinV-, 7AAD^-^) AML blast cells, calculated from six measurements for each condition. Statistical analysis was performed using paired T test (***, P < 0.0005). (C) HL60 and U937 cells were cocultured with BMSC WT/KO in medium containing exosome-depleted FCS (10%) for one month. Surviving AML cells maintained on WT or KO BMSC were collected for FACS analysis. Histogram represents EEF1A2 protein levels normalized to levels in AML cells maintained on BMSC WT ± SEM calculated from the analysis of 3 independent experiments. Statistical analysis was performed using the two-way analysis of variance (ANOVA) (**P < 0.005; ***P < 0.0005) (D) Flow cytometry analysis of AKT, pS473-AKT, pT450-AKT, RPS6 and pS235/236-RPS6 levels in HL60 cells, exposed for 1 month to WT versus KO BMSCs. Histogram represents protein levels normalized to levels in HL60 cells maintained on BMSC WT± SEM calculated from the analysis of 3 independent experiments. Statistical analysis was performed using the two-way analysis of variance (ANOVA) (*P < 0.05; **P < 0.005; ***P < 0.0005).

We also tested for the effects of long-term co-culture, using HL60 and U937 human AML cell lines (**Figure 6C**). In agreement with the data obtained for FLB1 in syntenin-deficient animals, EEF1A2 expression was significantly higher in HL60 and U937 cells co-cultured with KO BMSC (**Figure 6C & S5A**). This was associated with higher panAKT levels and an elevated phosphorylation of both pS473AKT and pS235/236-S6, further mimicking *in vivo* results obtained with FLB1 (**Figure 6D & S5B**).

In summary, syntenin-deficient BMSC support the survival and the education of several models and types of AML and suffice for that effect, suggesting a major role of the syntenin-null stroma in AML reprogramming.

### Sustained AML survival relies on BMSC autophagy

Next, we investigated whether recapitulation of the gain of AML aggressiveness (i.e. gain of cell survival) observed *in vitro* in the presence of KO BMSC requires close cell contacts. Strikingly, the use of transwells in the co-cultures strongly attenuated the increase of AML survival noted in the presence of KO BMSC (**Figure S6A**). Yet, we still noticed a slight increase of AML survival with conditioned media (CM) isolated from KO BMSC, and that effect was more marked compared to the effect of CM from WT BMSC (**Figure S6B**). Syntenin is implicated in exosome formation, and syntenin-deficient BMSC produce exosomes that are less loaded with typical exosomal cargo such as CD63 (**Figure S6C**). The effect on cell survival persisted when CM were depleted of exosomes by ultracentrifugation (**Figure S6B**). Altogether, these results suggest the importance of direct cell-to-cell contact for AML survival or, alternatively, “secretory” exchanges, possibly non receptor-mediated exchanges, requiring high concentrations of solute or particles and close contact with the blasts.

Several studies provided evidence that autophagy in stromal cells promotes cancer cell growth/survival^32^. Of note, BMSC characterization revealed a significantly higher endoglin expression in syntenin-null BMSC (**Figure S4B & 4C**). This TGFβ co-receptor is known to mediate autophagy by regulating Beclin-1 expression^33^. Moreover, in glioma cells syntenin- knockdown increases the expression of autophagic markers^20^. We therefore examined the possible contribution of stromal autophagy to AML cell survival. Under basal conditions, expression levels of the autophagy related proteins ATG3, eIF5A, LC3B-I and LC3B-II were significantly increased in KO BMSC cells (**Figure 7A & 7B**). These levels were equal to or even exceeding the expressions of these markers in BMSC co-cultured with AML cells for 48h, coculture leading to marked and, at least for eIF5A and ATG3, significant increases of the cellular levels of these markers, at least in WT BMSC (**Figure 7A & 7C**). Consistent with these results, immunofluorescence analysis revealed large numbers of LC3-positive perinuclear vesicles in KO BMSC (and an increase of these numbers compared to WT BMSC), co-cultured with HL60 cells (**Figure 7D**) or with different other models of AML (**Figure S7A**). Finally, to investigate the implication of BMSC autophagy in the gain of AML survival, we treated BMSC with Chloroquine (CQ) or Bafilomycin A1 (BFA1) before co-culture with AML cells. BFA1 and CQ pretreatments significantly inhibited HL60 survival on BMSC KO, by 59 and 22% respectively. Of note, on WT BMSC only BFA1 treatment affected HL60 survival, by 27% (**Figure 7E**). Similar results were obtained with the U937 model (**Figure S7C**). In summary, these data indicate that AML cells better survive in contact with syntenin-deficient stromal cells by exploiting autophagy pathways initiated and constitutively activated in the syntenin-deficient microenvironment.

**Figure 7.**
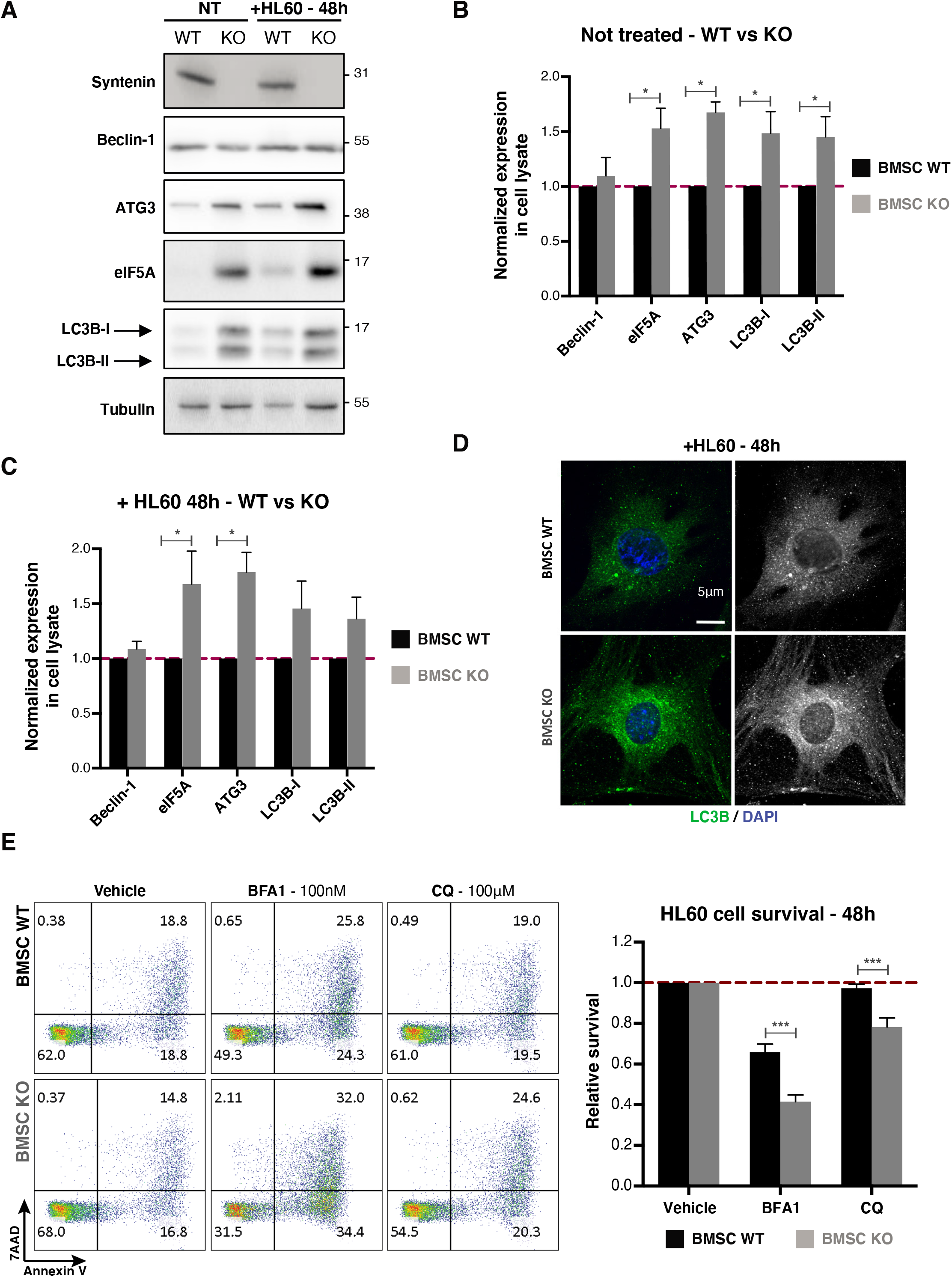
Increased autophagy in syntenin-deficient BMSC supports AML cell survival. (A) Total cell lysates from BMSC WT/KO in basal conditions (NT) or co-cultured with HL60 cells for 48 hours, analyzed by western blot for different autophagic markers, as indicated. (B) Histograms represent mean signal intensities of the indicated autophagic markers ± SEM in cell lysates, relative to signals obtained at basal condition in BMSC WT, calculated from the analysis of 9 independent experiments. Statistical analysis was performed using the two-way analysis of variance (ANOVA) (*P < 0.05). (C) Histograms represent mean signal intensities of the indicated autophagic markers ± SEM in cell lysates, relative to signals obtained in BMSC WT cocultured with HL60 at 48h, calculated from the analysis of 9 independent experiments. Statistical analysis was performed using the two-way analysis of variance (ANOVA) (*P < 0.05). (D) Representative immunofluorescent imaging of LC3B (green) and DAPI nuclear stain (blue), comparing WT and KO BMSCs, co-cultured with HL60 cells. (E) BMSC WT/KO were pre-treated for 6h with vehicle (Ctrl), Bafilomycin A1 (BFA1; 100nM) or Chloroquine (CQ; 100μM). Pretreated BMSC were washed extensively to remove excess of CQ or BFA1, HL60 cells were added and then both cell types were co-cultured in medium containing exosome-depleted FCS (10%). After 48h of co-culture, AML cells were collected and stained for apoptosis markers. Histograms represent mean of cell survival ± SEM relative to signal obtained with vehicle for each condition (BMSC WT or KO) and calculated from the analysis of 4 independent experiments. Statistical analysis was performed using the two-way analysis of variance (ANOVA) (***P < 0.0005).

## Discussion

Tumor-stroma interactions play a prominent role in the evolution of cancers. In contrast with the role of host-syntenin in melanoma progression^22^, we here provide evidence that leukemic blasts confronted with a syntenin-negative environment acquire a cell-autonomous advantage and aggressiveness.

Specifically, our proteomic analysis indicates a marked upregulation of EEF1A2, a factor shown to have pro-oncogenic activities in various cancers. EEF1A2 was originally described as a translation factor implicated in the delivery of aminoacyl-tRNAs to the A site of the ribosome^34^. EEF1A2 has pro-oncogenic activity by enhancing cancer cell proliferation and invasion, inhibiting apoptosis and regulating oxidative stress^34^. Preclinical data have described anti- eEF1A (narciclasine, mycalamide B) disrupting translation and protein synthesis as viable strategies to combat cancers^35,36^. Several clinical studies associated elevated EEF1A2 levels with poor prognosis^37,38^. As to hematological malignancies, it was reported that EEF1A2 expression is increased in multiple myeloma cell lines^28^ and that targeting EEF1A2 has antiproliferative effect in K562 cells, an erythro/myeloid leukemia cell line^39^. Xiao and colleagues have recently demonstrated that EEF1A2 is over-expressed in multiple AML cell lines, promoting cell proliferation and migration, but mainly decreasing apoptosis^40^. Here too, the upregulation of EEF1A2 appears notably linked to the reduction of apoptosis characterizing aggressive FLB1. Pharmacological inhibition of EEF1A2 with metarrestin (ML-246) indeed suggests that EEF1A2 drives the acquisition of a cell-autonomous survival, through the activation of Akt/RPS6 signaling pathway. While metarrestin is currently in phase I clinical trial for subjects with metastatic solid tumors, our data support the notion that such treatment could be extended to patients with AML.

Since previous studies highlighted the importance of post-transcriptional dysregulation in AML progression^41^, both transcriptional and epigenetic regulations/modifications of EEF1A2 (and also other up/down regulated factors) might be at work here. Indeed, it was recently reported that dimethylation of EEF1A at lysine 55 plays a key role in the EEF1A2-mediated oncogenesis of AML^40^. EEF1A2 appears to be also regulated by several onco-suppressor miRNAs^42^. Dysregulation of miRNAs have been found in multiple hematological malignancies^43^. Conceivably, miRNA and miRNA exchanges may induce stable phenotypic changes in cells.

Several studies have indicated BMSC can provide survival and anti-apoptotic signals to AML cells and as a result support leukemogenesis^44^,^45^. Consistent with such evidence, we here demonstrate that syntenin-deficient BMSC are able and suffice to educate different types of AML, enhancing cell survival and recapitulating at least some of the micro-environmental effects on AML observed *in vivo.* A deeper understanding of the mechanisms by which the BMSC are protecting/reprogramming AML blasts may contribute to the development of novel therapeutic approaches. Exosomes are potentially of specific interest in this respect, as vehicles of important mediators (proteins, lipids and nucleic acids, including miRNAs) of intercellular communication. In contrast with the variety of studies on BMSC remodeling by leukemia-derived exosomes^46^, reports on the relationship between BMSC-derived exosomes and leukemogenesis remain relatively rare. Yet, exosomes derived from cells in the tumor environment contribute to the behavior and evolution of tumor cells^47^. Loss of syntenin has well established negative effects on exosome biogenesis. Our findings highlight the importance of close contact between BMSC and blasts for the gain in AML survival. Thus, direct cell-to-cell contact, or secreted factors needed at high concentration (i.e. close to the source of production) seem to be involved in the gain of AML survival. Ultra-centrifugation, depleting the conditioned media of particles, however, seems to rather exclude a role for BMSC-derived exosomes in the gain of AML survival supported by these media. Altogether, these results suggested that syntenin-knockout alters still another cellular process implicated in intercellular communication.

Autophagy is an important homeostatic mechanism responsible for the elimination of damaged and abounding macromolecules such as proteins, lipids or damaged organelles. It also enables the recycling of nutrients, amino acids, and lipids and sustains organelle functions. Of note, so potentially also does the degradation of endosomal intraluminal vesicles and the secretion, (re)uptake and lysosomal degradation of exosomes and exosome-cargo, exchanges that at least in part are lost when syntenin is missing. Recent reports also identified a role for autophagy in unconventional protein secretion. Autophagy is increasingly recognized as an important pathway in cancer initiation, maintenance and resistance to therapy^32^. High levels of basal autophagy correlate with a unique secretory profile when compared to cancer cells with lower levels of autophagy^48^. Several recent reports also highlighted how autophagy induced in stromal cells that make up the tumor microenvironment promotes tumor cell survival, growth and invasion^49,50^. Enhanced basal autophagy in cancer-associated fibroblasts (CAFs) facilitates the secretion of cytokines (IL6, IL8) and promotes head and neck cancer progression^51^. Secretory autophagy also controls the secretion of metabolites, such as alanine, by microenvironment cells of pancreatic cancer, supporting PDAC growth under nutrient-limiting conditions^52^. Mitophagy, the degradation of damaged mitochondria, has been recently associated with tumor stroma crosstalk. AML cells are prone to accept mitochondria from BMSC, increasing oxidative phosphorylation and resistance to chemotherapy^53^. Yet our proteomic data are not supportive of such process in the present context. Fisher’s group has identified syntenin as regulator of protective autophagy in glioblastoma cells, by triggering FAK/PKC/BCL2 and EGFR signaling (suppressing high levels of autophagy elicited by cell detachment from the extracellular matrix and promoting resistance to anoikis)^20^. Our findings identify a ‘constitutive’ dysregulation of autophagy in syntenin-deficient BMSC. In basal condition or in presence of AML cells, KO-BMSC show an important increase of the autophagy related proteins eIF5A and ATG3. This is associated with elevated levels of LC3B lipidation in basal condition, and an increase of LC3B-positives vesicles. These proteins are known to regulate autophagy by controlling autophagosome formation. eIF5A indeed mediates the translation of ATG3 protein that is required for lipidation of LC3B protein^54^. Intriguingly, eIF5A was shown to regulate p53 activity in collaboration with syntenin: when eIF5A interacts with syntenin, the eIF5A-induced increase in p53 protein level is significantly inhibited^55^. Like Sox4, eIF5A interacts with the N-terminal domain of syntenin. Syntenin-binding protects Sox4 from proteasomal degradation, affecting Sox4 transcriptional output^56^. Yet, as demonstrated by the sequestration of GSK regulating beta-catenin, also the biology of the multivesicular body (i.e. intraluminal vesicle formation) is involved in the stabilization of cytosolic proteins, regulating transcriptional output^57^. It remains to be explored whether syntenin might affect eIF5A accumulation in cells by similar mechanisms, but the data strongly support the notion that syntenin acts as a repressor of eIF5A activities, thus leading to autophagy dysregulation in syntenin-knockout BMSC. Finally, pharmacological inhibitors demonstrate that autophagy induced by syntenin depletion in the stroma participates to AML cell survival. It remains to be established, however, whether autophagy induced by syntenin deficiency enhances the release of soluble factors/cytokines, metabolites or both.

In summary, we show that a syntenin-deficient microenvironment supports leukemic cell expansion, reprogramming blasts and ultimately leading to the activation of cell autonomous survival mechanisms in these cells. A dysregulation of autophagy in the syntenin-deficient microenvironment is at least in part responsible for the gain of AML cell survival. Syntenin is potentially an interesting target for tumor therapy^58,59^. Unless the importance of cell autonomous effects of syntenin loss/inhibition prevails for tumor development, current observations, however, question the suitability or safety of syntenin as a systemic target in cancer therapy and AML in particular.

## Acknowledgements

We thank Prof. Olivier Herault (University of Tours, France) for providing us with FLB1 cells, Prof. Peter Carmeliet (University of Leuven, Belgium) for providing us with FLPeR mice and Prof. Pierre Close and his team (University of Liège, Belgium) for advices with protein synthesis *in vivo* experiments. This work was supported by grants from the Concerted Actions Program of the KU Leuven (GOA/12/016), the National Research Agency (ANR, Investissements d’Avenir, A*MIDEX project ANR-11-IDEX-0001-02), the Fund for Scientific Research-Flanders (Fonds Wetenschappelijk Onderzoek—Vlaanderen Grants G.0846.15 and G0C5718N), the Institut National du Cancer (INCa, subvention 2013-105), and the Belgian Foundation against cancer (STK, FA/2014/294). RL was the recipient of fellowships from the ARC French Foundation for Cancer Research, Inserm and Institut Paoli-Calmettes.

## Authorship and conflict-of-interest statements

The experiments were designed by RL, JF and RC; RL, JF, AG, RC, LC, MB, RG, BB and CF performed the experiments; RL, PZ, RC and RC analyzed the data; MAL, YC, NV, JPB contributed new reagents/analytic tools and provide advices; and RL, GD, and PZ conceived the project, supervised the work and wrote the paper. All authors were invited to adjust the manuscript.

## Conflict-of-interest disclosure

The authors declare no competing financial interests.

## Notes

### Competing Interest Statement

The authors have declared no competing interest.

## References

1. Döhner H, Estey E, Grimwade D, et al. Diagnosis and management of AML in adults: 2017 ELN recommendations from an international expert panel. Blood. 2017;

2. Ayala F, Dewar R, Kieran M, Kalluri R. Contribution of bone microenvironment to leukemogenesis and leukemia progression. Leukemia. 2009;

3. Shiozawa Y, Taichman RS. Dysfunctional niches as a root of hematopoietic malignancy. Cell Stem Cell. 2010;

4. Ghobrial IM, Detappe A, Anderson KC, Steensma DP. The bone-marrow niche in MDS and MGUS: Implications for AML and MM. Nat. Rev. Clin. Oncol. 2018;

5. Rashidi A, DiPersio JF. Targeting the leukemia-stroma interaction in acute myeloid leukemia: rationale and latest evidence. Ther. Adv. Hematol. 2016;

6. Ding L, Zhang W, Yang L, et al. Targeting the autophagy in bone marrow stromal cells overcomes resistance to vorinostat in chronic lymphocytic leukemia. Onco. Targets. Ther.2018;

7. Grootjans JJ, Zimmermann P, Reekmans G, et al. Syntenin, a PDZ protein that binds syndecan cytoplasmic domains. Proc. Natl. Acad. Sci. U. S. A. 1997;

8. Zimmermann P, Tomatis D, Rosas M, et al. Characterization of syntenin, a syndecan- binding PDZ protein, as a component of cell adhesion sites and microfilaments. Mol. Biol. Cell. 2001;

9. Beekman JM, Coffer PJ. The ins and outs of syntenin, a multifunctional intracellular adaptor protein. J. Cell Sci. 2008;

10. Friand V, David G, Zimmermann P. Syntenin and syndecan in the biogenesis of exosomes. Biol. Cell. 2015;

11. Shimada T, Yasuda S, Sugiura H, Yamagata K. Syntenin: PDZ protein regulating signaling pathways and cellular functions. Int. J. Mol. Sci. 2019;

12. Baietti MF, Zhang Z, Mortier E, et al. Syndecan-syntenin-ALIX regulates the biogenesis of exosomes. Nat. Cell Biol. 2012;

13. Ghossoub R, Lembo F, Rubio A, et al. Syntenin-ALIX exosome biogenesis and budding into multivesicular bodies are controlled by ARF6 and PLD2. Nat. Commun. 2014;

14. Kashyap R. Syntenin and syndecan in the biogenesis of exosomes. Sci. Rep. 2020;in press:

15. Rabouille C. KRS: A cut away from release in exosomes. J. Cell Biol. 2017;

16. Chen J, Tang J, Chen W, et al. Effects of syndecan-1 on the expression of syntenin and the migration of U251 glioma cells. Oncol. Lett. 2017;

17. Oyesanya RA, Bhatia S, Menezes ME, et al. MDA-9/Syntenin regulates differentiation and angiogenesis programs in head and neck squamous cell carcinoma. Oncoscience. 2014;

18. Das SK, Sarkar D, Emdad L, Fisher PB. MDA-9/Syntenin: An emerging global molecular target regulating cancer invasion and metastasis. Adv. Cancer Res. 2019;

19. Talukdar S, Das SK, Pradhan AK, et al. Novel function of MDA-9/Syntenin (SDCBP) as a regulator of survival and stemness in glioma stem cells. Oncotarget. 2016;

20. Talukdar S, Pradhan AK, Bhoopathi P, et al. MDA-9/Syntenin regulates protective autophagy in anoikis-resistant glioma stem cells. Proc. Natl. Acad. Sci. U. S. A. 2018;

21. Yu Y, Li S, Wang K, Wan X. A PDZ Protein MDA-9/Syntenin: As a Target for Cancer Therapy. Comput. Struct. Biotechnol. J. 2019;

22. Das SK, Guo C, Pradhan AK, et al. Knockout of MDA-9/Syntenin (SDCBP) expression in the microenvironment dampens tumor-supporting inflammation and inhibits melanoma metastasis. Oncotarget. 2016;

23. Wilhelm BT, Briau M, Austin P, et al. RNA-seq analysis of 2 closely related leukemia clones that differ in their self-renewal capacity. Blood. 2011;

24. Farley FW, Soriano P, Steffen LS, Dymecki SM. Widespread recombinase expression using FLPeR (Flipper) mice. genesis. 2000;

25. Signer RAJ, Magee JA, Salic A, Morrison SJ. Haematopoietic stem cells require a highly regulated protein synthesis rate. Nature. 2014;

26. Pellegrino R, Calvisi DF, Neumann O, et al. EEF1A2 inactivates p53 by way of PI3K/AKT/mTOR-dependent stabilization of MDM4 in hepatocellular carcinoma. Hepatology. 2014;

27. Sun Y, Du C, Wang B, et al. Up-regulation of eEF1A2 promotes proliferation and inhibits apoptosis in prostate cancer. Biochem. Biophys. Res. Commun. 2014;

28. Li Z, Qi CF, Shin DM, et al. Eef1a2 promotes cell growth, inhibits apoptosis and activates JAK/STAT and AKT signaling in mouse plasmacytomas. PLoS One. 2010;

29. Frankowski KJ, Wang C, Patnaik S, et al. Metarrestin, a perinucleolar compartment inhibitor, effectively suppresses metastasis. Sci. Transl. Med. 2018;

30. Brenner AK, Nepstad I, Bruserud Ø. Mesenchymal stem cells support survival and proliferation of primary human acute myeloid leukemia cells through heterogeneous molecular mechanisms. Front. Immunol. 2017;

31. Corradi G, Baldazzi C, Očadlíková D, et al. Mesenchymal stromal cells from myelodysplastic and acute myeloid leukemia patients display in vitro reduced proliferative potential and similar capacity to support leukemia cell survival. Stem Cell Res. Ther. 2018;

32. Folkerts H, Hilgendorf S, Vellenga E, Bremer E, Wiersma VR. The multifaceted role of autophagy in cancer and the microenvironment. Med. Res. Rev. 2019;

33. Pan CC, Kumar S, Shah N, et al. Endoglin regulation of Smad2 function mediates beclin1 expression and endothelial autophagy. J. Biol. Chem. 2015;

34. Abbas W, Kumar A, Herbein G. The eEF1A proteins: At the crossroads of oncogenesis, apoptosis, and viral infections. Front. Oncol. 2015;

35. Van Goietsenoven G, Hutton J, Becker JP, et al. Targeting of eEF1A with Amaryllidaceae isocarbostyrils as a strategy to combat melanomas. FASEBJ. 2010;

36. Dang Y, Schneider-Poetsch T, Eyler DE, et al. Inhibition of eukaryotic translation elongation by the antitumor natural product Mycalamide B. RNA. 2011;

37. Kawamura M, Endo C, Sakurada A, et al. The prognostic significance of eukaryotic elongation factor 1 alpha-2 in non-small cell lung cancer. Anticancer Res. 2014;

38. Giudici F, Petracci E, Nanni O, et al. Elevated levels of eEF1A2 protein expression in triple negative breast cancer relate with poor prognosis. PLoS One. 2019;

39. Losada A, Muñoz-Alonso MJ, García C, et al. Translation elongation factor eEF1A2 is a novel anticancer target for the marine natural product plitidepsin. Sci. Rep. 2016;

40. Xiao S, Wang Y, Ma Y, et al. Dimethylation of eEF1A at Lysine 55 Plays a Key Role in the Regulation of eEF1A2 on Malignant Cell Functions of Acute Myeloid Leukemia. Technol. Cancer Res. Treat. 2020;

41. Wallace JA, O’Connell RM. MicroRNAs and acute myeloid leukemia: Therapeutic implications and emerging concepts. Blood. 2017;

42. Vislovukh A, Kratassiouk G, Porto E, et al. Proto-oncogenic isoform A2 of eukaryotic translation elongation factor eEF1 is a target of miR-663 and miR-744. Br. J. Cancer. 2013;

43. Liao Q, Wang B, Li X, Jiang G. miRNAs in acute myeloid leukemia. Oncotarget. 2017;

44. Garrido SM, Appelbaum FR, Willman CL, Banker DE. Acute myeloid leukemia cells are protected from spontaneous and drug-induced apoptosis by direct contact with a human bone marrow stromal cell line (HS-5). Exp. Hematol. 2001;

45. Forte D, García-Fernández M, Sánchez-Aguilera A, et al. Bone Marrow Mesenchymal Stem Cells Support Acute Myeloid Leukemia Bioenergetics and Enhance Antioxidant Defense and Escape from Chemotherapy. Cell Metab. 2020;

46. Kumar B, Garcia M, Murakami JL, Chen CC. Exosome-mediated microenvironment dysregulation in leukemia. Biochim. Biophys. Acta - Mol. Cell Res. 2016;

47. Hoshino A, Costa-Silva B, Shen TL, et al. Tumour exosome integrins determine organotropic metastasis. Nature. 2015;

48. Kraya AA, Piao S, Xu X, et al. Identification of secreted proteins that reflect autophagy dynamics within tumor cells. Autophagy. 2015;

49. Katheder NS, Khezri R, O’Farrell F, et al. Microenvironmental autophagy promotes tumour growth. Nature. 2017;

50. Mowers EE, Sharifi MN, Macleod KF. Functions of autophagy in the tumor microenvironment and cancer metastasis. FEBSJ. 2018;

51. New J, Arnold L, Ananth M, et al. Secretory autophagy in cancer-associated fibroblasts promotes head and neck cancer progression and offers a novel therapeutic target. Cancer Res. 2017;

52. Sousa CM, Biancur DE, Wang X, et al. Pancreatic stellate cells support tumour metabolism through autophagic alanine secretion. Nature. 2016;

53. Moschoi R, Imbert V, Nebout M, et al. Protective mitochondrial transfer from bone marrow stromal cells to acute myeloid leukemic cells during chemotherapy. Blood. 2016;

54. Lubas M, Harder LM, Kumsta C, et al. eIF 5A is required for autophagy by mediating ATG 3 translation. EMBO Rep. 2018;

55. Li AL, Li HY, Jin BF, et al. A novel eIF5A complex functions as a regulator of p53 and p53- dependent apoptosis. J. Biol. Chem. 2004;

56. Beekman JM, Vervoort SJ, Dekkers F, et al. Syntenin-mediated regulation of Sox4 proteasomal degradation modulates transcriptional output. Oncogene. 2012;

57. Taelman VF, Dobrowolski R, Plouhinec JL, et al. Wnt signaling requires sequestration of Glycogen Synthase Kinase 3 inside multivesicular endosomes. Cell. 2010;

58. Kegelman TP, Wu B, Das SK, et al. Inhibition of radiation-induced glioblastoma invasion by genetic and pharmacological targeting of MDA-9/Syntenin. Proc. Natl. Acad. Sci. U. S. A. 2017;

59. Leblanc R. Pharmacological inhibition of syntenin PDZ2 domain impairs breast cancer cell activities and exosome loading with syndecan and EpCAM cargo. J. Extracell. vesicles. 2020;

