## supplemental text for "A syntenin-deficient microenvironment educates AML for aggressiveness"

### Supplementary data

#### Supplementary methods

##### Mass Spectrometry Analysis and Protein Quantification.

*Sample preparation and labelling***.** Equal amounts of protein (i.e. around 100µg) were aliquoted and adjusted to 100 µl using 100 mM TEAB. Proteins were reduced using TCEP and then alkylated using iodoacetamide as mentioned in the TMT mass tag labelling instructions (Thermo Scientific). Proteins were precipitated using six volumes of cold acetone at -20°C overnight. The pellet was then suspended in 100 mM TEAB and digested with 2.5µg Trypsin (overnight at 37°C, Promega). Peptides were labelled with TMT reagents as described in the TMT labelling instructions (Thermo Scientific). Briefly, samples were transferred to TMT reagents reconstituted using 40ul of anhydrous acetonitrile and then incubated at room temperature for one hour. Five biological replicates for both conditions, i.e. PWT and PKO FLB1, were distributed in the ten TMT channels. The reaction was quenched by adding 8 µl of 5% hydroxylamine and incubating for 15 min. Labelled peptides were mixed, and dried to remove organic solvent prior to clean-up via Sep-Pak (50 mg C18 SepPak; Waters). Labeled peptide mixtures were separated in 80 fractions via high-pH reversed phase chromatography (Waters; Xbridge C18 analytical column) and concatenated into thirteen fractions. Samples were dried and stored at −80 °C prior to LC-MS analysis.

*Mass spectrometry analysis*. Samples were reconstituted in 4% acetonitrile/0.1% trifluoroacetic acid prior to being analysed by liquid chromatography (LC)-tandem mass spectrometry (MS/MS) with an Orbitrap Fusion Lumos Tribrid Mass Spectrometer (Thermo Fischer) online with an Ultimate 3000RSLCnano chromatography system

(Dionex). Peptides were separated on a reverse phase LC EASY-Spray C18 column from Dionex (PepMap RSLC C18, 50 cm x 75 µm I.D, 100 Å pore size, 2 µm particle size) at 300 nL/min flow rate and 40°C and using a two steps linear gradient (4-20% acetonitrile/H20; 0.1 % formic acid for 180 min and 20-45-45% acetonitrile/H2O; 0.1 % formic acid for 60 min. An EASY-Spray nanosource was used for peptide ionization (2 200 V, 275°C). The mass spectrometer was used in data dependent mode to switch consistently between MS1, MS2 for peptide identification and multinotch-MS3 for protein abundance measurements. Time between Masters Scans was set to 3 seconds. MS1 spectra were acquired with the Orbitrap in the range of m/z 375-1500 at a FWHM resolution of 120 000 measured at 200 m/z. AGC target was set at 4.0x10E5 with a 50 ms maximum injection time. The more abundant precursor ions were selected for MS2 (Top speed 3 seconds) and collision induced dissociation fragmentation at 35% was performed and analysed in the linear ion trap using the "Inject Ions for All Available Parallelizable time" option with a maximum injection time of 105 ms and an AGC target of 1.0x10E5. For Multi-Notch MS3, the top 10 precursor ions from each MS2 scan were fragmented by HCD followed by orbitrap analysis (Resolution = 60 000; First mass = 100 m/z; AGQ target = 250000; MaxIT = 86ms, 100-500 m/z). Charge state screening was enabled to include precursors with 2 and 7 charge states. Dynamic exclusion was enabled with a repeat count of 1 and a duration of 60s. The mass spectrometry proteomics data have been deposited to the ProteomeXchange Consortium via the PRIDE partner repository with the data set identifier PXD (under submission).

*Database search and relative quantification.* Relative TMT intensity-based quantification was processed using the freely available MaxQuant computational proteomics platform, version 1.6.3.4. Raw data were searched against the mouse database, extracted from UniProt on the 9th of December 2019 and containing 20379 entries (reviewed) supplemented with known contaminants. Searches were performed with default parameters trypsin enzymatic cleavage with up to two missed cleavages, and three variable modifications allowed per peptide. Searches were performed with variable methionine oxidation (+15.99491), static cysteine carboxyamidomethylation (+57.02146), and static TMT modifications on lysine and the peptide N-termini (+229.16293). The false discovery rate (FDR) at the peptide and protein levels were set to 1% and determined by searching a reverse database. For protein grouping, all proteins that cannot be distinguished based on their identified peptides were assembled into a single entry according to the MaxQuant rules. TMT reporter intensities were corrected for isotope impurities according to the lot used. The statistical analysis was done with Perseus program (version 1.6.2.1). First corrected TMT reporter intensities were base 2 logarithmized to obtain a normal distribution and normalized by subtracting the median for each TMT channels. The few proteins with missing values were removed from the dataset. To determine whether a given detected protein was specifically differential, a two-sample t-test was done using permutation-based FDR-controlled at 0.01 and employing 250 permutations. The p value was adjusted using a scaling factor s0 with a value of 0.4 for the FLB1-P4WT/-P4KO with n=5.

**Western blots.** The proteins were heat-denaturated in Laemmli sample buffer, fractionated in 12.5% or 15% gels by SDS–PAGE and electro-transferred to nitrocellulose membrane. Membranes were stained with Ponceau red and immunoblotted with the indicated primary antibodies (**Table S1**), and HRP-conjugated secondary antibodies (Mouse or Rabbit, Thermofisher scientific; 1/10000). Signals were visualized using Amersham ECL Prime Western Blotting Detection Reagent (GE Healthcare).

**Isolation and culture of murine bone marrow stromal cells.** 8 weeks old female C57Bl/6J WT or KO mice were sacrificed, hind limbs were collected and stripped of their skin and muscle manually. For isolation of the primary cells, the bone marrow from the femora and the tibia was flushed in RPMI buffer containing 1% FCS and 1% of penicillin-streptomycin (Gibco, Carlsbad, CA, USA). Red blood cells were removed using ACK lysis buffer (Gibco, Carlsbad, CA, USA) 10min at room temperature. Cells were then washed twice into RPMI medium and seeded in a rat tail collagen-1 (5µg/cm$^2$; Gibco, Carlsbad, CA, USA) coated flask for cell culture. BMSC were routinely grown in RPMI medium supplemented with 10% FCS, 10% horse serum (Gibco, Carlsbad, CA, USA), 1%Penicillin-Streptomycin, and 1%Sodium-Pyruvate, in plastic flasks coated with collagen. The growing cells were characterized as bone marrow stromal cells by flow cytometry analysis: dead cells were excluded using DRAQ7 (Beckman Coulter, Brea, CA, USA)) and cells were stained with antibodies for the positive (Sca1, CD51, CD106, CD54, CD44, CD105 & CD295) and the negative (CD45, CD3e, CD11c, B220, CD11b, CD19 & Ter119) selection of this lineage. BMSC WT/KO were used at the fourth to the tenth passages and always irradiated at 30Gy 24h prior co-culture experiments.

***In vitro* differentiation.** Osteoblastic differentiation was induced by culturing the BMSC WT/KO for 4 weeks with 50 µg ml−1 l-ascorbic acid 2-phosphate, 10 mM glycerophosphate (Sigma-aldrich, Saint-Louis, MO, USA) and 15% FBS in α-MEM with penicillin-streptomycin. Adipocyte differentiation was induced with 1 µM dexamethasone, 10 µg ml−1 insulin (Sigma-Aldrich, Saint-Louis, MO, USA) and 10% FBS in α-MEM with penicillin-streptomycin. All cultures were maintained with 5% CO2 in a water-jacketed incubator at 37 °C, and media change was performed every 2-3 days. To assess *in vitro* differentiation into mesenchymal lineages, cells were briefly washed with phosphate buffered saline solution (PBS1X; Gibco, Carlsbad, CA, USA) and fixed for 10 min at room temperature with 4% paraformaldehyde (Santa Cruz). For the detection of mineral calcium deposition, osteoblasts were stained with Alizarin Red solution (Sigma-Aldrich, Saint-Louis, MO, USA) for 30min and rinsed 3 times with distillated water. Adipocytes were stained with Oil Red O (Sigma-Aldrich, Saint-Louis, MO, USA) as follows: cells were washed with 60% isopropanol and allowed to dry completely. Oil Red O working solution was prepared as a 6:4 dilution in distilled water of a 0.35 g ml−1 Oil Red O solution in isopropanol (Sigma-aldrich, Saint-Louis, MO, USA) and filtered 20 min later. Cells were incubated for 60 min with Oil red O working solution and rinsed four times using distilled water.

***In vivo* homing assays.** For homing assays, a total of $3\times10^6$ cells/mouse was injected into the retro-orbital vein of WT or KO C57Bl/6J mice. 16 hours after cell inoculation, animals were sacrificed, bone marrow, spleen and lymph nodes were collected for analysis. Number of FLB1 in the bone marrow and the spleen were evaluated by flow cytometry by using the CD45.1-FITC antibody and CountBright™ Absolute Counting Beads according to the manufacturer.

**In vivo limiting dilution.** For extreme limiting dilution assays (EDLA), five doses (20, 200, 500, 1000, 2000) of CD45.1+ cells, isolated from WT or KO animals at graft 4, were transplanted into C57Bl/6J mice. The cut-off for FLB1 engraftment was the exhibition of more than 10% of CD45.1+ cells in the PB.

**Measurement of protein synthesis.** For *in vivo* analysis, OP-Puro (50 mg $kg^{-1}$ body mass; pH 6.4–6.6 in PBS1X; Medchem Source, Federal Way, WA, USA) was injected intraperitoneally. One hour later mice were euthanized, BM was collected, and $3 \times 10^6$ cells were stained with CD45.1 antibody at room temperature for 30min. Cells were then washed twice in $Ca^{2+}$- and $Mg^{2+}$-free PBS1X and fixed in 0.5 ml of 1% paraformaldehyde in PBS1X for 15 min on ice. Cells were washed in PBS, then permeabilized in 200 µl PBS supplemented with 3% fetal bovine serum and 0.1% saponin (Sigma-aldrich, Saint-Louis, MO, USA) for 5 min at room temperature (20–25 °C). The azide-alkyne cycloaddition was performed using the Click-iT Cell Reaction Buffer Kit (Life Technologies, Carlsbad, CA, USA) and azide conjugated to Alexa Fluor 555 (Life Technologies, Carlsbad, CA, USA) at 5 µM final concentration. After the 30-min reaction, the cells were washed twice in PBS1X supplemented with 3% fetal bovine serum and 0.1% saponin, then resuspended in PBS1X supplemented with 4′,6-diamidino-2-phenylindole (DAPI; 4 µg $ml^{-1}$ final concentration) and analysed by flow cytometry. Relative rates of protein synthesis were calculated by normalizing OP-Puro signals to whole bone marrow after subtracting autofluorescence background. For *in vitro* analysis, 0.5 x $10^6$ cells were seeded in 2ml of RPMI medium containing exosome-depleted FCS (10%). OP-Puro (50 µM final concentration) was added to the culture medium for 1 h. The cells were collected, fixed, permeabilized, and the azide-alkyne cycloaddition was performed as described above. Mean OP-Puro fluorescence reflected absolute fluorescence values for each cell population from multiple independent experiments.

**Cell cycle assays.** Distribution of cell cycle phases was determined by Propidium Iodide (PI; Life Technologies, Carlsbad, CA, USA) staining and flow cytometry analysis. Briefly, 5 × $10^6$ BM cells were isolated and stained with CD45.1 antibody. Cells were fixed in 1 mL ice-cold 70% ethanol for 30min at 4°C and stored at −20 °C. At the time of analysis, the cells were centrifugated, washed once again in PBS1X and stained with a freshly made solution containing 40µ/ml PI, 0.1% Triton x-100 and 40µ/ml RNaseA (Life Technologies, Carlsbad, CA, USA) in PBS1X. Percentages of cells within cell cycle compartments (G1, S and G2/M) were determined by using FACS LSRII flow cytometer and data were analyzed with Flowjo software.

**Reverse transcription-polymerase chain reaction (RT-PCR).** RNA was extracted from cells using Nucleospin RNA isolation kit (Macherey-Nagel). Reverse Transcription was done using PrimeScript RT Reagent with gDNA Eraser kit (TAKARA CLONTECH) and MJ Research PTC-220 Dyad PCR System (Conquer Scientific). Quantitative PCR reactions were performed using SsoAdvanced Universal Sybr Green Supermix kit (BIORAD) with specific primers and analyzed using CFX96 Touch Real-Time PCR Detection System (BIORAD). Each PCR cycle consisted of 15s of denaturation at 95°C, 30s of annealing and extension at 60°C.

**Transwell assays.** BMSC WT/KO were co-cultured with AML cell lines. In order to differentiate the role of direct cell-cell interaction from the role of secreted components, co-culture experiments were performed in direct contact or using Transwells (0.4 μm pore size, polycarbonate membrane cell culture insert, Corning) to separate the cell populations. Passage 4 to 10 BMSC WT/KO were detached, counted and irradiated at 30Gy. A total of 10,000 irradiated BMSC were seeded in 12-well plates pre-coated with type-I collagen (5μg/$cm^2$; Gibco) and were cultured in complete medium for 24h. Then, 50,000 AML cells were added to each well and maintained in short-term co-culture for 24h to 96h.

**Exosomes and total cell lysates.** For comparative analyses, exosome-enriched fractions were collected from equivalent amounts of culture medium, conditioned by equivalent amounts of BMSC WT or KO. After 16hours, cell media were collected and exosomes were isolated by three sequential centrifugation steps at 4 °C: 10 min at 500×g, to remove cells; 30 min at 10,000×g, to remove cell debris; and 1h30min at 100,000×g, to pellet exosomes (exosome-enriched fraction), followed by one wash with 1400µL of PBS1X (100,000×g, 1h), to remove soluble serum and secreted proteins. Exosomal pellets were then re-suspended in 100µL of PBS1X. The lysates from corresponding cultures were cleared by centrifugation at 1500rpm for 5 min and then resuspended in lysis buffer (TrisHCL pH 7.4 30mM, NaCl 150mM, 1% NP40 (IGEPAL), 1µg/ml aprotinin, 1µg/ml leupeptin). Although only little variations were observed from sample to sample, exosomal amounts loaded for the western blot were normalized according to the number of parent cells from where exosomes were secreted.

**Nanoparticle tracking analysis (NTA).** Concentrations and size distributions of exosomal fractions (100.000 x g pellet) isolated from the same number of cells were measured at a dilution of 1:50 in 500 µl of PBS1 with the Nanosight NS300 instrument (Malvern), which was equipped with a 488 nm laser and a sCMOS camera. Three videos of 60 sec each were recorded for each sample at 25°C and -used to calculate mean values of particle concentration.

**Immunofluorescence staining and confocal microscopy.** Passage 4 to 10 BMSC WT/KO were detached, counted and irradiated at 30Gy. A total of 2,000 irradiated BMSC were seeded in collagen type-I (5µg/cm$^2$; Gibco) pre-coated 8-well permanox chamber slides (Lab-Tek) and were cultured in complete medium for 24h. Then, 10,000 AML cells were added to each well and maintained in short-term co-culture. After 48h, AML cells were removed, BMSC were fixed with 100% Methanol for 5 min, washed in PBS and permeabilized with 0.1% Triton X-100 for 5 min. Cells were saturated with PBS buffer containing 1% BSA, 10% normal goat serum, 0.3M glycine and in 0.1% Triton for 1h. Cells were then incubated with the indicated antibodies in saturation buffer. Cells were mounted in ProLong™ Diamond Antifade Mountant containing DAPI and observed with a

Zeiss confocal microscope (LSM 880, Zeiss, France) with the corresponding lasers and 40× objectives. Confocal images were analyzed using Photoshop (Adobe, San Jose, CA) software.

**Supplementary figure legends**

**Figure S1. FLB1 homing in WT & KO animals, FLB1 invasion of the bone marrow and the spleen upon serial transplantations and serial back-transplantations.** (A) Non-irradiated 8-11wks old wild-type (WT) and syntenin-deficient (KO) mice were injected with 50,000 FLB1 cells in the retro-orbital vein. Leukemia progression was monitored weekly by FACS analysis using the CD45.1-FITC antibody for detection of the FLB1 cells in the peripheral blood (PB). At day 21, animals the PB, the bone marrow (BM) and the spleen were collected for further analysis (Left panel). The dot-plot summarizes the percentage of CD45.1$^+$ cells ± SEM, in the PB, the BM and the spleen of WT and KO mice, 21 days after cell inoculation, as measured by flow cytometry. Each symbol represents an individual animal. Statistical analysis was performed using the nonparametric Mann-Whitney U test (Right panel). (B) Non-irradiated 8-10wks old WT and KO mice were injected with $3x10^6$ FLB1 cells in the retro-orbital vein. 12 hours after cell inoculation, animals were sacrificed and BM, spleen and lymph nodes were collected for analysis (Left panel). Number of FLB1 in the BM and the spleen were evaluated by FACS, using FITC-conjugated CD45.1-antibody. Results are expressed as mean of the number of CD45.1$^+$ cells per organ, ± SEM. Statistical analysis was performed using the nonparametric Mann-Whitney U test. (C) WT and KO mice were injected with FLB1 cells derived from consecutive (1-4) serial engraftments in the corresponding host, and were sacrificed at the survival threshold, i.e. when blast levels reached >10% in the PB. Summary of the CD45.1$^+$ cell frequencies measured in the BM and the spleen of engrafted WT and KO mice, at the survival threshold. Percentages and mean percentage of CD45.1+ cells, ± SEM. Statistical analysis was performed using the nonparametric Mann-Whitney U test. (D) FLB1 maintained for up to four consecutive passages in KO mice, and controls maintained in WT mice, were serially further transplanted in WT mice, for up to three passages. All animals were sacrificed at the survival threshold, i.e. when blast levels reached >10% in the PB. CD45.1+ cell frequencies, and means ± SEM, measured in the BM and the spleen of the mice, at the survival threshold. Statistical analysis was performed using the nonparametric Mann-Whitney U test.

**Figure S2. Syntenin-deficiency of the host does not affect the homing, the cell cycle or the clonogenicity of AML cells.** (A) Non-irradiated 8-10wks old WT and KO mice were, respectively, injected with $3x10^6$ FLB1-P4WT or FLB1-P4KO cells in the retro-orbital vein. 12 hours after cell inoculation, animals were sacrificed and BM, spleen and lymph nodes were collected for analysis. (Left panel). Numbers of FLB1 cells in the BM and the spleen were counted by FACS, using the CD45.1-FITC antibody and CountBright™ Absolute Counting Beads according to the manufacturer. Results are expressed as mean number of $CD45.1^+$ cells per organ ± SEM. Statistical analysis was performed using the nonparametric Mann-Whitney U test. (B) FLB1-P4WT cells were injected at day 0 into WT background, while the inoculation of FLB1-P4KO cells into KO mice (evolving more rapidly) was delayed for one week. Cell cycle was analyzed in both early and late phases of leukemia progression. At day 14 & 21, animals were sacrificed; BM was collected, fixed and stained with propidium iodide. The cell cycle stage of the $CD45.1^+$ cells was then analyzed by FACS (left panel). Results are expressed as percentage of cells in the different phases of the cell cycle ± SEM. Statistical analysis was performed using the nonparametric Mann-Whitney U test (right panel). (C) Extreme limiting dilution assay performed by injecting decreasing numbers of sorted FLB1-P4WT or FLB1-P4KO cells. The number of mice ultimately demonstrating the presence of at least 10% of CD45.1+ cells in the peripheral blood, as well as the number of transplanted mice is indicated for each cell dose. Leukemic initiating cells (LIC) frequency, as well as the upper and lower estimates for each sample are reported (Left Panel). Log-fraction plot of the extreme limiting dilution model (EDLA) fitted to FLB1-P4WT vs FLB1-P4KO. The slope of the line is the log-active cell fraction. The dotted lines indicate the 95% confidence intervals. Stem cell frequencies of FLB1-P4WT (black lines) and FLB1-P4KO (red lines) are respectively 1/401 and 1/365 (Right panel).

**Figure S3. OP-Puro analysis.** Gating on viable cells, OP-Puro labeling was combined with staining with anti-CD45.1 to enable separate quantification of protein synthesis in AML cells and nonmalignant hematopoietic cells. Flow cytometry gating strategy of OP-Puro incorporation in both $CD45.1^-$ and $CD45.1^+$ cells isolated from both WT and KO animals, 1 h after OP-Puro administration *in vivo*.

**Figure S4. BMSC isolation & characterization.** (A) 5-6wks old WT and KO mice were sacrificed, hind limbs were collected, and bones were flushed. Whole BM was plated on collagen coated flasks and expanded *ex vivo*. Hematopoietic cells were gradually eliminated by several washes, medium changes and at least four passages. (B) Murine BMSC WT/KO at passage 4 after isolation were analyzed by flow cytometry, for the expression of the indicated BMSC markers. (C) Real-time PCR analysis of indicated marker mRNA expressions in BMSC, WT versus KO. Values were normalized to housekeeping L32/GAPDH genes. Data represent the mean ± SEM of 5 independent experiments performed in duplicate. Statistical analysis was performed using unpaired T test (**, $P < 0.005$; ***, $P < 0.0005$). (D) BMSCs at passage 4 were cultivated under specific osteogenic or adipogenic conditions for 21 days. Alizarin Red and Red oil staining were performed to assess, respectively, Ca2++ deposition after osteogenic induction and lipid droplet accumulation after adipocyte differentiation.

**Figure S5. AML/BMSC long term co-cultures.** (A) Flow cytometry analysis of EEF1A2 level in HL60 and U937 cells, exposed for 1 month to WT versus KO BMSCs. (B) Flow cytometry analysis of AKT, pS473-AKT, pT450-AKT, RPS6 & pS236/236-RPS6 levels in HL60 cells, after 1 month of co-culture with BMSC, WT or KO.

**Figure S6. Relevance of stromal syntenin exosomal pathways for AML cell survival.** (A) FLB1 cells were co-cultured, in direct contact or in transwells, with BMSC WT/KO. After 48h, blasts were collected and stained for apoptosis markers. Results are expressed as mean percentage of living (AnnexinV$^-$, 7AAD$^-$) CD45.1$^+$ cells ± SEM calculated for 3 independent experiments. Statistical analysis was performed using the one-way analysis of variance (ANOVA) (*$P < 0.05$; **$P < 0.01$). (B) Media conditioned by WT or by KO BMSC (for 48h) were depleted or not depleted of exosomes by ultracentrifugation (100,000xg, 1h at 4°C). FLB1 cells treated for 48h with conditioned medium were stained for apoptosis markers. Results are expressed as mean percentages of living (AnnexinV$^-$ , 7AAD$^-$) cells. (C) BMSC, WT or KO, were cultured in medium containing exosome-depleted FCS (10%) for 16h. Conditioned medium was submitted to differential centrifugation and particles pelleting at 100,000 x g were analyzed by western blotting (*Left panel*). Total cell lysates and corresponding pelleted extracellular particles were analyzed by western blot, tracing several exosomal markers, as indicated (*Right panel*). (D)

**Figure S7. BMSC WT/KO autophagy in co-culture with AML cells.** (A) Representative immunofluorescent imaging of LC3B (green) and DAPI nuclear stain (blue), comparing WT and KO BMSCs, co-cultured with FLB1 or U937 cells. (C) BMSC WT/KO were pre-treated for 6h with vehicle (Ctrl), Bafilomycin A1 (BFA1; 100nM) or Chloroquine (CQ;

100μM). Pre-treated BMSC were washed extensively to remove excess of CQ or BFA1, U937 cells were added and then both cell types were co-cultured in medium containing exosome-depleted FCS (10%). After 48h of co-culture, AML cells were collected and stained for apoptosis markers. Histograms represent mean of cell survival ± SEM relative to signal obtained with vehicle for each condition (BMSC WT or KO) from one experiment performed in triplicate. Statistical analysis was performed using the two-way analysis of variance (ANOVA) (**$P < 0.005$).

**Table S1. Antibodies and reagents**

**Table S2. FLB1P4WT and FLB1P4KO proteomic dataset related to Figure 2A**.

**Table S3. Diseases and functions Ingenuity analysis of differentially expressed proteins.**

**Table S4**. **AML patient characteristics versus blast cell survival.** Survival data are expressed as mean percentages of living (AnnexinV$^-$ 7AAD$^-$) blast cells, calculated from six measurements for each condition. Statistical analysis was performed using 2-way analysis of variance (ANOVA) (***, $P < 0.001$).
