## Supplementary material for "A syntenin-deficient microenvironment educates AML for aggressiveness": supp figures

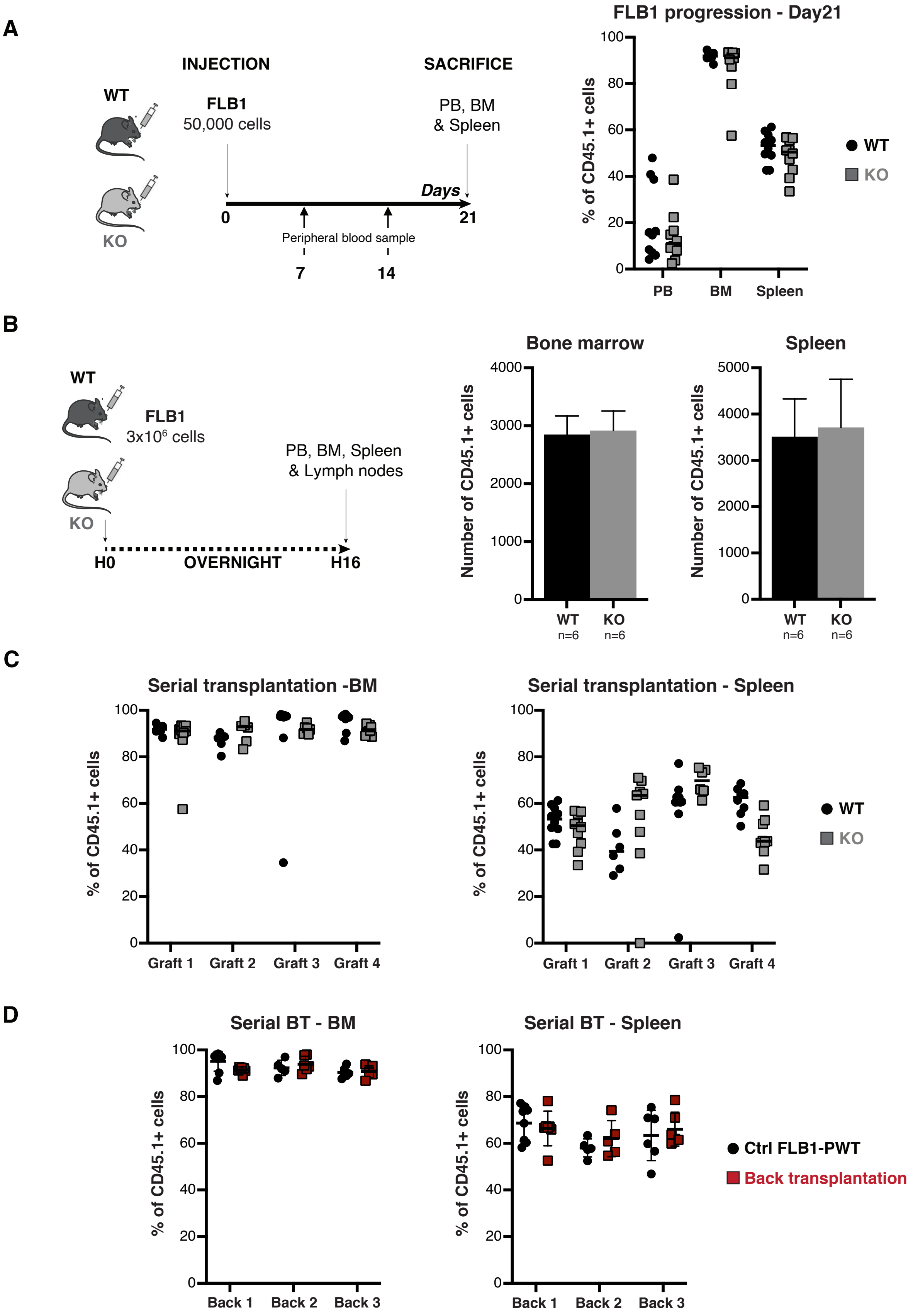

A
INJECTION
SACRIFICE
WT
KO
FLB1
50,000 cells
PB, BM
& Spleen
Days
0
21
Peripheral blood sample
7
14
FLB1 progression - Day21
% of CD45.1+ cells
PB
BM
Spleen
WT
KO
B
WT
KO
FLB1
3x10^6 cells
PB, BM, Spleen
& Lymph nodes
H0
OVERNIGHT
H16
Bone marrow
Spleen
Number of CD45.1+ cells
WT
n=6
KO
n=6
C
Serial transplantation -BM
Serial transplantation - Spleen
% of CD45.1+ cells
Graft 1
Graft 2
Graft 3
Graft 4
WT
KO
D
Serial BT - BM
Serial BT - Spleen
% of CD45.1+ cells
Back 1
Back 2
Back 3
Ctrl FLB1-PWT
Back transplantation


Figure S1

**A**

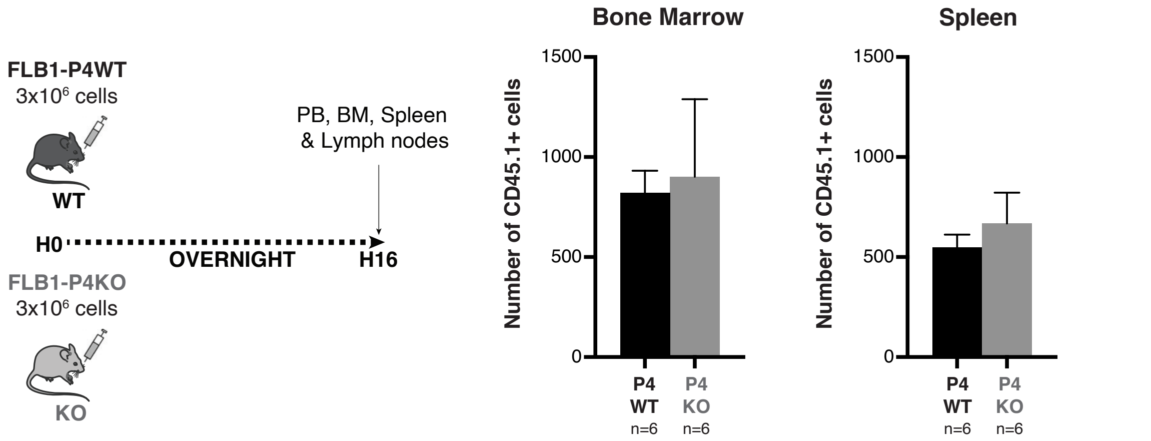


**B**

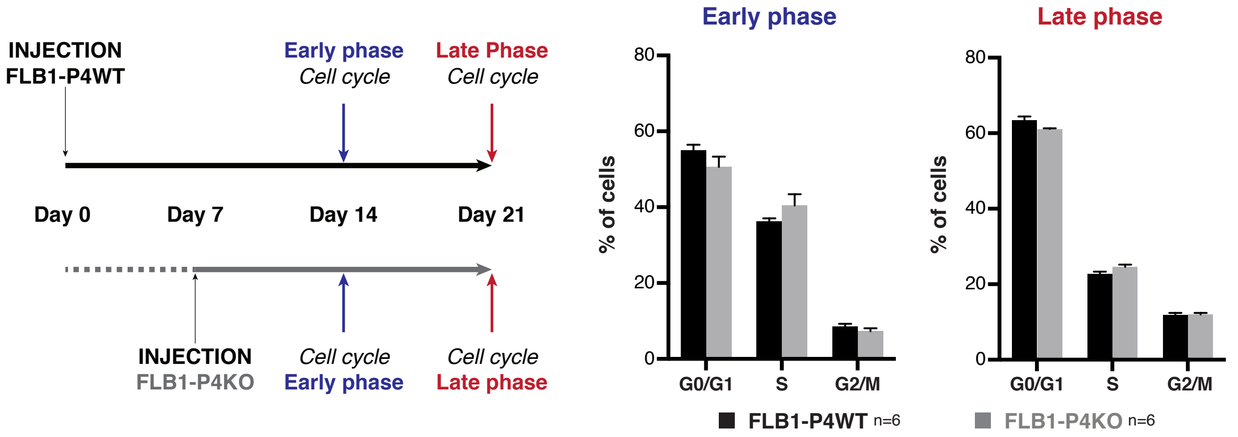


**C**

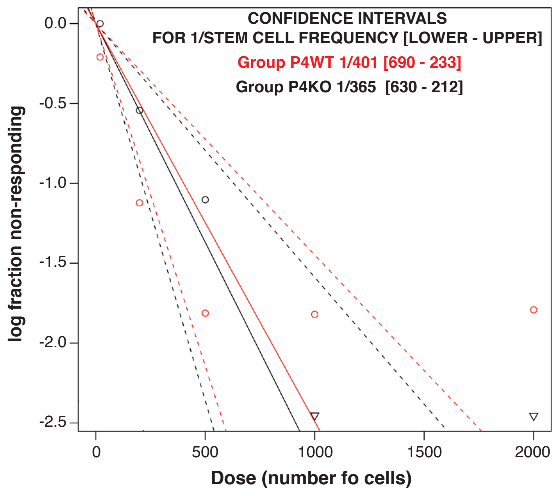


| Group | Dose (cells) | Mice tested | Mice engrafted (%of CD45.1>10% in PB) | Confidence intervals for 1/stem cell frequency [lower - upper] |
|---|---|---|---|---|
| FLB1-P4WT | 2000 | 6 | 5 | 1/401 [691 - 233] |
| | 1000 | 6 | 5 | |
| | 500 | 6 | 5 | |
| | 200 | 12 | 8 | |
| | 20 | 6 | 1 | |
| FLB1-P4KO | 2000 | 6 | 6 | 1/365 [630 - 212] |
| | 1000 | 6 | 6 | |
| | 500 | 6 | 4 | |
| | 200 | 12 | 5 | |
| | 20 | 6 | 0 | |

**Figure S2**

A

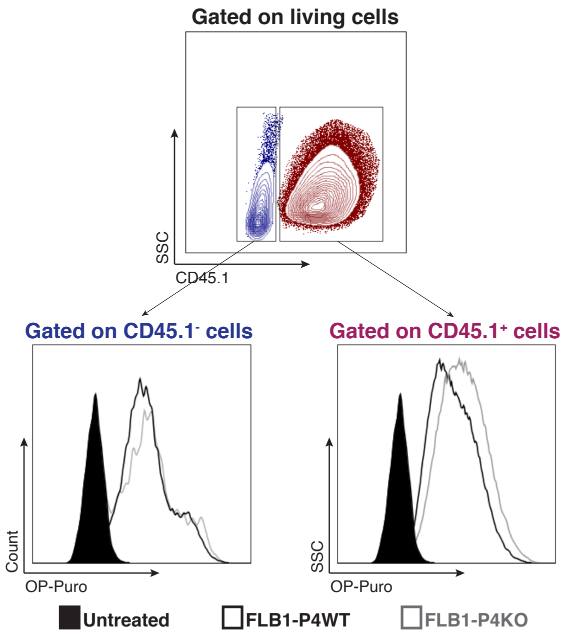
Gated on living cells
SSC
CD45.1
Gated on CD45.1⁻ cells
Gated on CD45.1⁺ cells
Count
OP-Puro
SSC
OP-Puro
Untreated
FLB1-P4WT
FLB1-P4KO

Figure S3

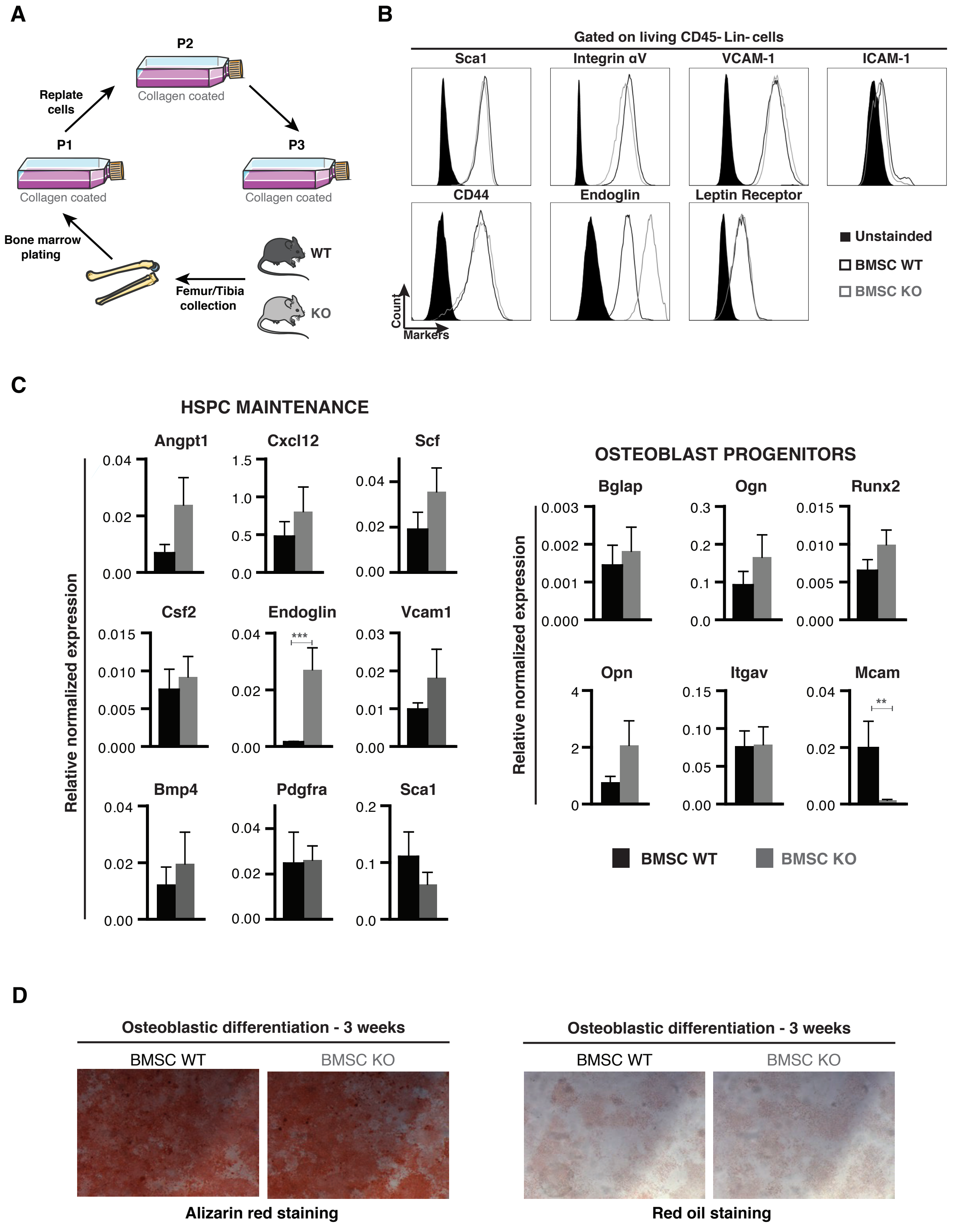
A
P2
Replate cells
Collagen coated
P1
P3
Bone marrow plating
WT
Femur/Tibia collection
KO
B
Gated on living CD45- Lin- cells
Sca1
Integrin αV
VCAM-1
ICAM-1
CD44
Endoglin
Leptin Receptor
Count
Markers
Unstainded
BMSC WT
BMSC KO
C
HSPC MAINTENANCE
Angpt1
Cxcl12
Scf
Csf2
Endoglin
Vcam1
Bmp4
Pdgfra
Sca1
Relative normalized expression
OSTEOBLAST PROGENITORS
Bglap
Ogn
Runx2
Opn
Itgav
Mcam
BMSC WT
BMSC KO
D
Osteoblastic differentiation - 3 weeks
BMSC WT
BMSC KO
Alizarin red staining
Red oil staining

Figure S4

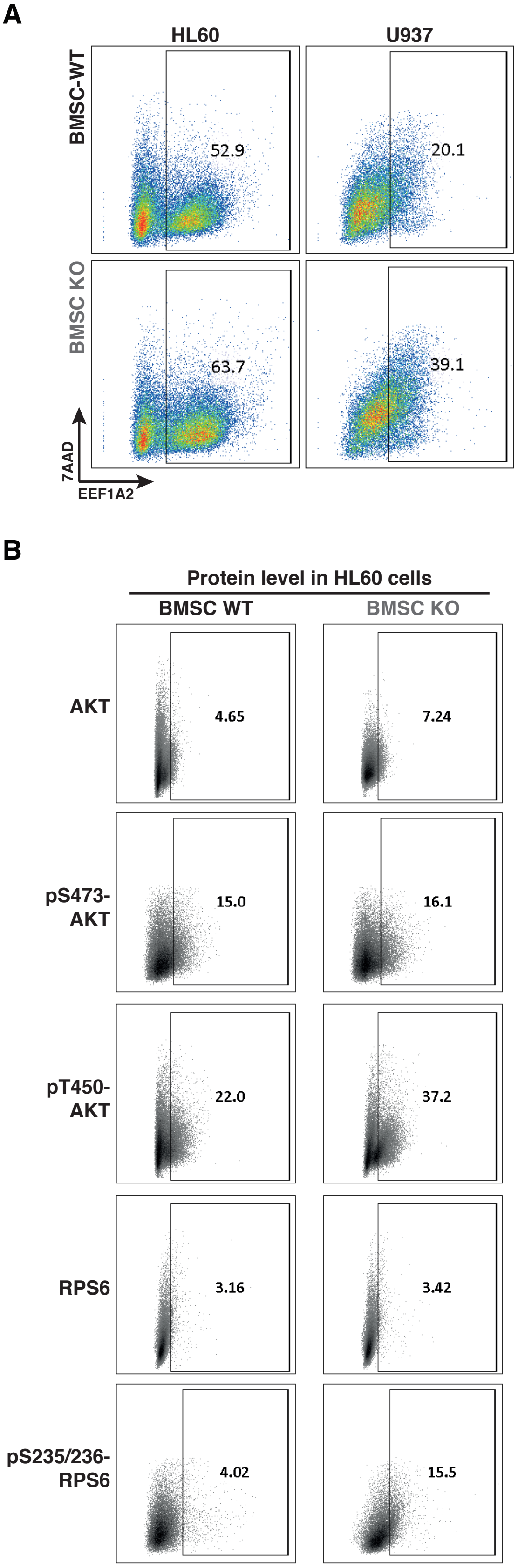
A
HL60
U937
BMSC-WT
BMSC KO
52.9
20.1
63.7
39.1
7AAD
EEF1A2
B
Protein level in HL60 cells
BMSC WT
BMSC KO
AKT
4.65
7.24
pS473-AKT
15.0
16.1
pT450-AKT
22.0
37.2
RPS6
3.16
3.42
pS235/236-RPS6
4.02
15.5

**Figure S5**

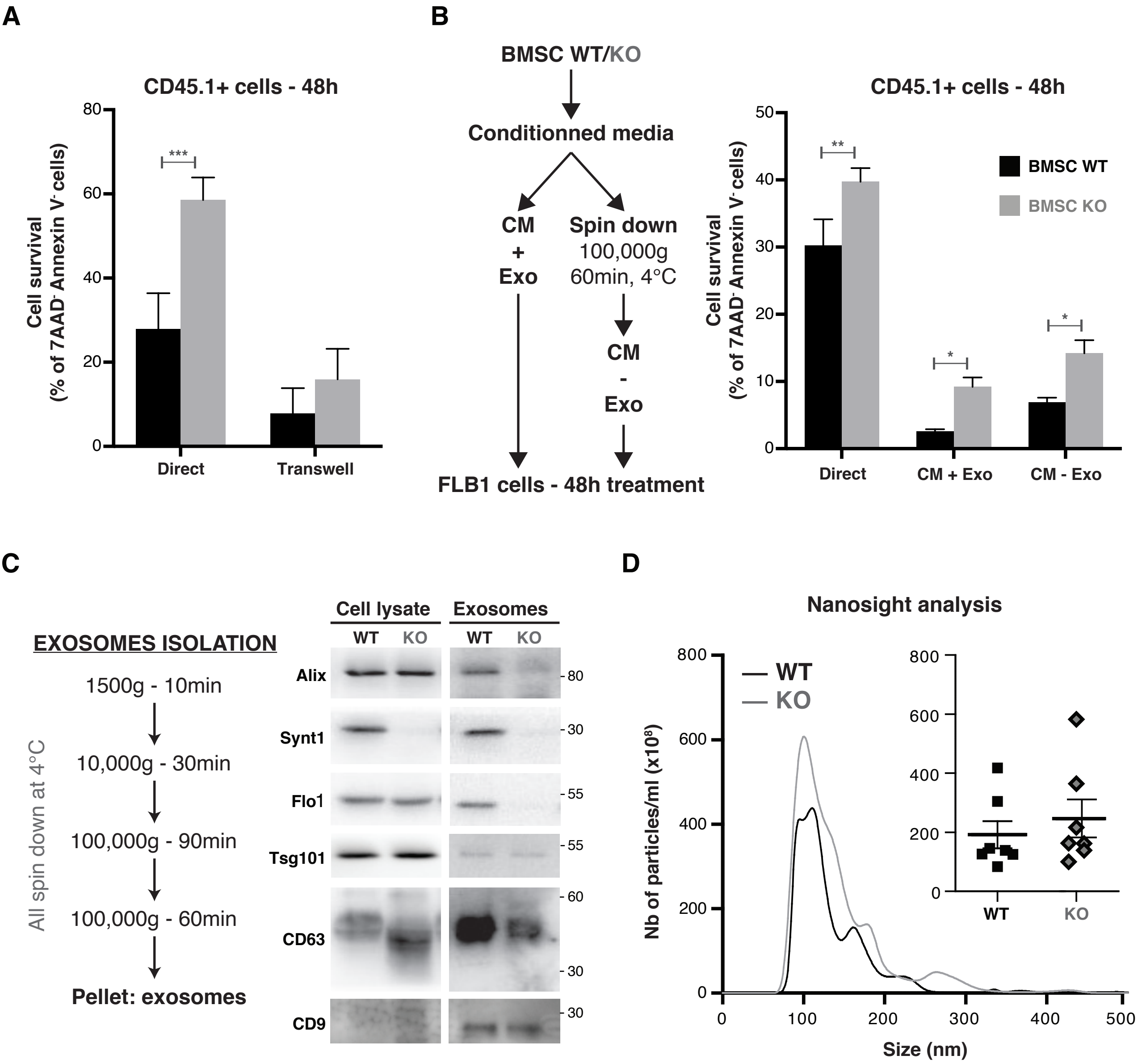
A
CD45.1+ cells - 48h
Cell survival (% of 7AAD- Annexin V- cells)
***
Direct
Transwell
B
BMSC WT/KO
Conditionned media
CM + Exo
Spin down 100,000g 60min, 4°C
CM - Exo
FLB1 cells - 48h treatment
CD45.1+ cells - 48h
BMSC WT
BMSC KO
**
*
*
Direct
CM + Exo
CM - Exo
C
EXOSOMES ISOLATION
1500g - 10min
10,000g - 30min
100,000g - 90min
100,000g - 60min
Pellet: exosomes
All spin down at 4°C
Cell lysate
Exosomes
WT
KO
Alix
Synt1
Flo1
Tsg101
CD63
CD9
D
Nanosight analysis
WT
KO
Nb of particles/ml (x10^8)
Size (nm)

Figure S6

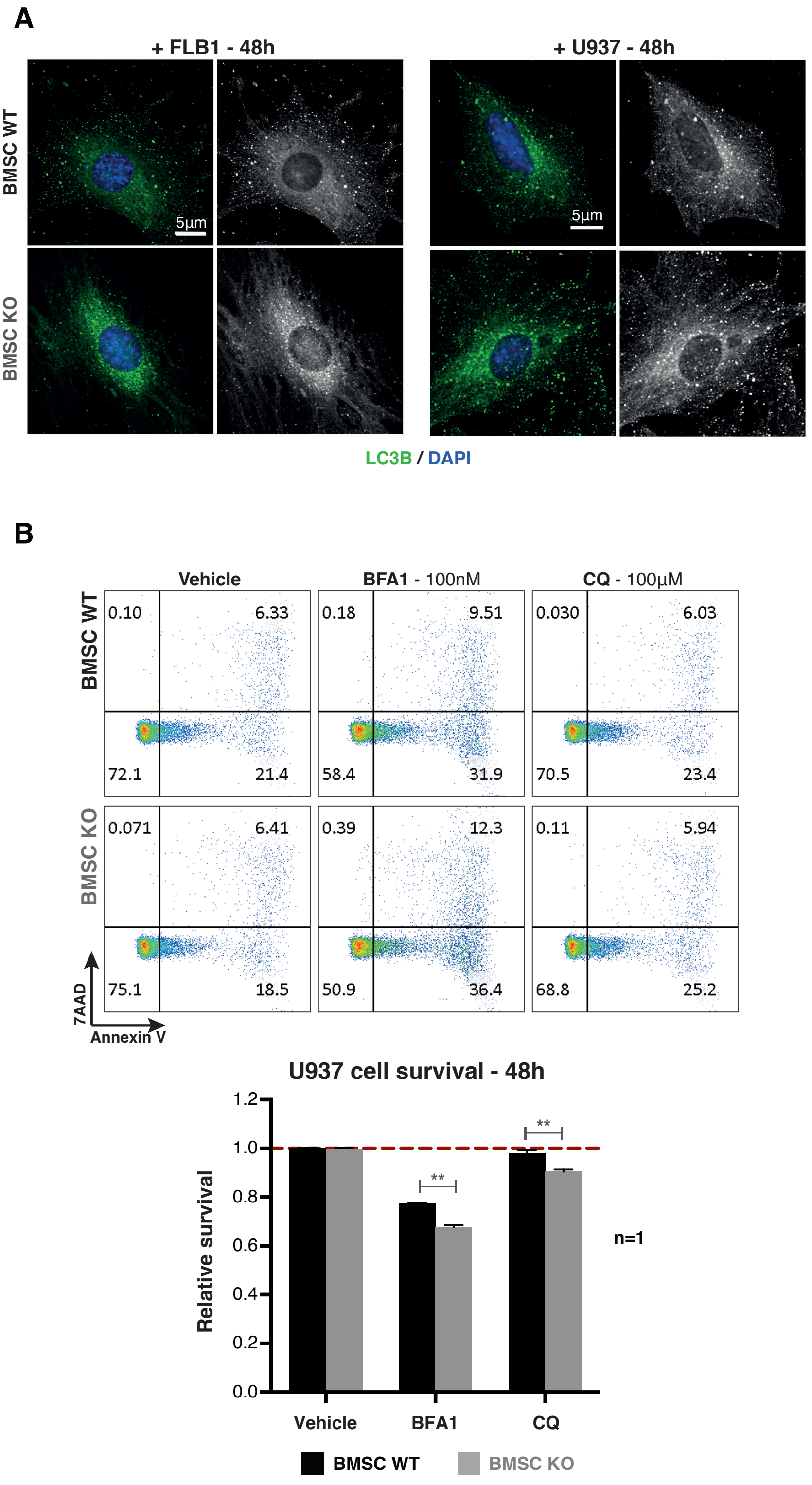

A
+ FLB1 - 48h
+ U937 - 48h
BMSC WT
BMSC KO
5µm
5µm
LC3B / DAPI
B
Vehicle
BFA1 - 100nM
CQ - 100µM
BMSC WT
BMSC KO
0.10
6.33
72.1
21.4
0.18
9.51
58.4
31.9
0.030
6.03
70.5
23.4
0.071
6.41
75.1
18.5
0.39
12.3
50.9
36.4
0.11
5.94
68.8
25.2
7AAD
Annexin V
U937 cell survival - 48h
Relative survival
1.2
1.0
0.8
0.6
0.4
0.2
0.0
**
**
Vehicle
BFA1
CQ
n=1
BMSC WT
BMSC KO


**Figure S7**
